# IFT88 transports Gucy2d, a guanylyl cyclase, to maintain sensory cilia function in *Drosophila*

**DOI:** 10.1101/2020.12.15.417840

**Authors:** Sascha Werner, Sihem Zitouni, Pilar Okenve-Ramos, Susana Mendonça, Anje Sporbert, Christian Spalthoff, Martin C. Göpfert, Swadhin Chandra Jana, Mónica Bettencourt-Dias

**Author notes:** Correspondence should be addressed to Sascha Werner; Swadhin Chandra Jana; Mónica Bettencourt-Dias. Shared lead authors (Swadhin Chandra Jana, Mónica Bettencourt-Dias).

## Abstract

Cilia are involved in a plethora of motility and sensory-related functions. Ciliary defects cause several ciliopathies, some of which with late-onset, suggesting cilia are actively maintained. While much is known about cilia assembly, little is understood about the mechanisms of their maintenance. Given that intraflagellar transport (IFT) is essential for cilium assembly, we investigated the role of one of its main players, IFT88, in ciliary maintenance. We show that DmIFT88, the *Drosophila melanogaste*r orthologue of IFT88, continues to move along fully formed sensory cilia, and that its acute knockdown in the ciliated neurons of the adult affects sensory behaviour. We further identify DmGucy2d, the *Drosophila* guanylyl cyclase 2d, as a DmIFT88 cargo, whose loss also leads to defects in sensory behaviour maintenance. DmIFT88 binds to the intracellular part of DmGucy2d, which is evolutionarily conserved and mutated in several degenerative retina diseases, taking the cyclase into the cilia. Our results offer a novel mechanism for the maintenance of sensory cilia function and its potential role in human diseases.

## Introduction

Cilia are microtubule (MT)-based organelles that emanate from the surface of eukaryotic cells and are vital for several functions, including motility and sensing (Breslow and Holland, 2019; Reiter and Leroux, 2017; Sreekumar and Norris, 2019). Cilia biogenesis is a multistep process that is tightly coordinated with cell differentiation. A cilium consists of two regions: a ciliary base, and a ciliary protrusion or shaft. The latter grows from the ciliary base and is composed of a MT-based skeleton (the axoneme) and a ciliary membrane. Ciliary proteins are produced in the cell body and then move either by diffusion or by active transport through the ciliary base into the ciliary shaft. This active transport is called intraflagellar transport (IFT), which depends on molecular motors moving from the ciliary base to the axoneme tip (anterograde) and in reverse (retrograde) direction (reviewed in (Breslow and Holland, 2019)).

Defects in cilia structure and function cause several human disorders, collectively called ciliopathies, which show manifestations such as alterations in body symmetry, obesity and cystic kidneys. While defects in cilia assembly can lead to numerous diseases, they do not account for all symptoms of many cilia-related disorders, such as retinitis pigmentosa, Nephronophthesis and Alstrom syndrome, which show progressive tissue-degeneration in the patients (Reiter and Leroux, 2017; Sreekumar and Norris, 2019). Given that many ciliated cells involved in sensory reception, such as photoreceptors, ciliated sensory neurons and epithelial cells are long-lived, it is possible that breakdown of ciliary maintenance is causal to those diseases. IFT has been implicated in maintaining ciliary structural and functional properties such as flagella/cilia length, or localisation of signalling receptors, such as Opsin, TRPV and Guanylyl Cyclases, into the cilium, in various organisms (Bhowmick et al., 2009; Eguether et al., 2014; Jiang et al., 2015; Marshall et al., 2005; van der Burght et al., 2020). In contrast, in the unicellular parasite *Trypanosoma*, IFT88 depletion does not affect the structure of fully formed flagella, only causing defects in flagella beating which suggests a deregulation of flagella signalling components (Fort et al., 2016). Furthermore, entry of several receptors, such as Smoothened (SMO) or SSTR3, into cilia is IFT-independent (Milenkovic et al., 2009; Williams et al., 2014; Ye et al., 2013). Altogether, the existing evidence suggests that the maintenance of ciliary structure and sensory function is a heterogeneous process, whose underlying mechanisms need to be explored.

Here we study the underpinnings of ciliary maintenance in a genetically tractable organism, *Drosophila melanogaster*. In particular, we focus on adult Type-I ciliated sensory neurons as they rarely get replenished (Fernández-Hernández et al., 2020; Grotewiel et al., 2005) and are involved in different sensory functions with straightforward experimental readouts, including hearing, climbing, gravitaxis and olfaction (Han et al., 2003; Jana et al., 2011). Given the importance of IFT in cilia assembly and maintenance in *Chlamydomonas* (Marshall et al., 2005; Pazour et al., 2002), we chose to investigate the function of a key IFT protein, IFT88, in the maintenance of *Drosophila* cilia structures and function. Here, we refer to the *Drosophila melanogaster* homologue of IFT88 protein as DmIFT88 (also known as NOMPB (Han et al., 2003)). We first found that DmIFT88 trains move within fully-grown chordotonal cilia, suggesting IFT plays an active role in *Drosophila* cilia maintenance. We next observed that depletion of DmIFT88 in adult ciliated sensory neurons leads to defects in climbing (or negative-gravitaxis) behaviour, while subtly affecting their structure. In the search for distinct DmIFT88 cargoes important for ciliary maintenance, we discovered that DmIFT88 binds and transports to the cilium the fly homologue of an evolutionarily conserved particulate guanylyl cyclase (Gucy2d) that is involved in human retina diseases. Our research shows that IFT is important for the functional maintenance of adult sensory cilia, in particular through transporting key signalling cargoes.

## Results

### IFT88 protein sequences show high conservation of structural domains

*Drosophila melanogaster* homologues of IFT proteins (i.e. A-complex, B-complex, and BBSome) have been identified through bioinformatics analysis (for a summary, see our compilation of all information in Supplemental Figure 1A, and Supplemental Table 1). Several of them were shown to be expressed in ciliated neurons or in the fly head (Avidor-Reiss et al., 2004; Chintapalli et al., 2007; Lee et al., 2008; Lee et al., 2018; Mourao et al., 2016), yet only a few, such as DmIFT88 and DmIFT140, have been mechanistically investigated (Han et al., 2003; Lee et al., 2008). Studies in other model organisms suggest that only a few evolutionarily conserved IFT proteins are critical for cilia formation (Eguether et al., 2014; Fan et al., 2010; Fort et al., 2016; Pazour et al., 2002). Among them is the IFT88 protein, and thus, we investigated whether it might also have an important role in transporting components needed for cilium maintenance. In *Drosophila melanogaster*, two isoforms of DmIFT88 have been reported, which are similar in their amino acid composition (Figure 1Ai; (Han et al., 2003)) and are predicted to bear 10 tetratricopeptide repeat (TPR) domains (Karpenahalli et al., 2007) (Figure 1Aii), which often act as interfaces for protein-protein interactions (Allan and Ratajczak, 2011; Taschner et al., 2012). In particular, all IFT88 homologues (in insects as well as vertebrates) are predicted to comprise 10 to 15 TPRs (Figure 1Aii), pointing to conserved vital roles of this protein in mediating protein interactions, either within the IFT-B core complex or between IFT and cargo (Taschner et al., 2012). We thus investigated whether DmIFT88 continues to transport cargoes after cilium assembly, which could be important for its maintenance.

**Figure 1:**
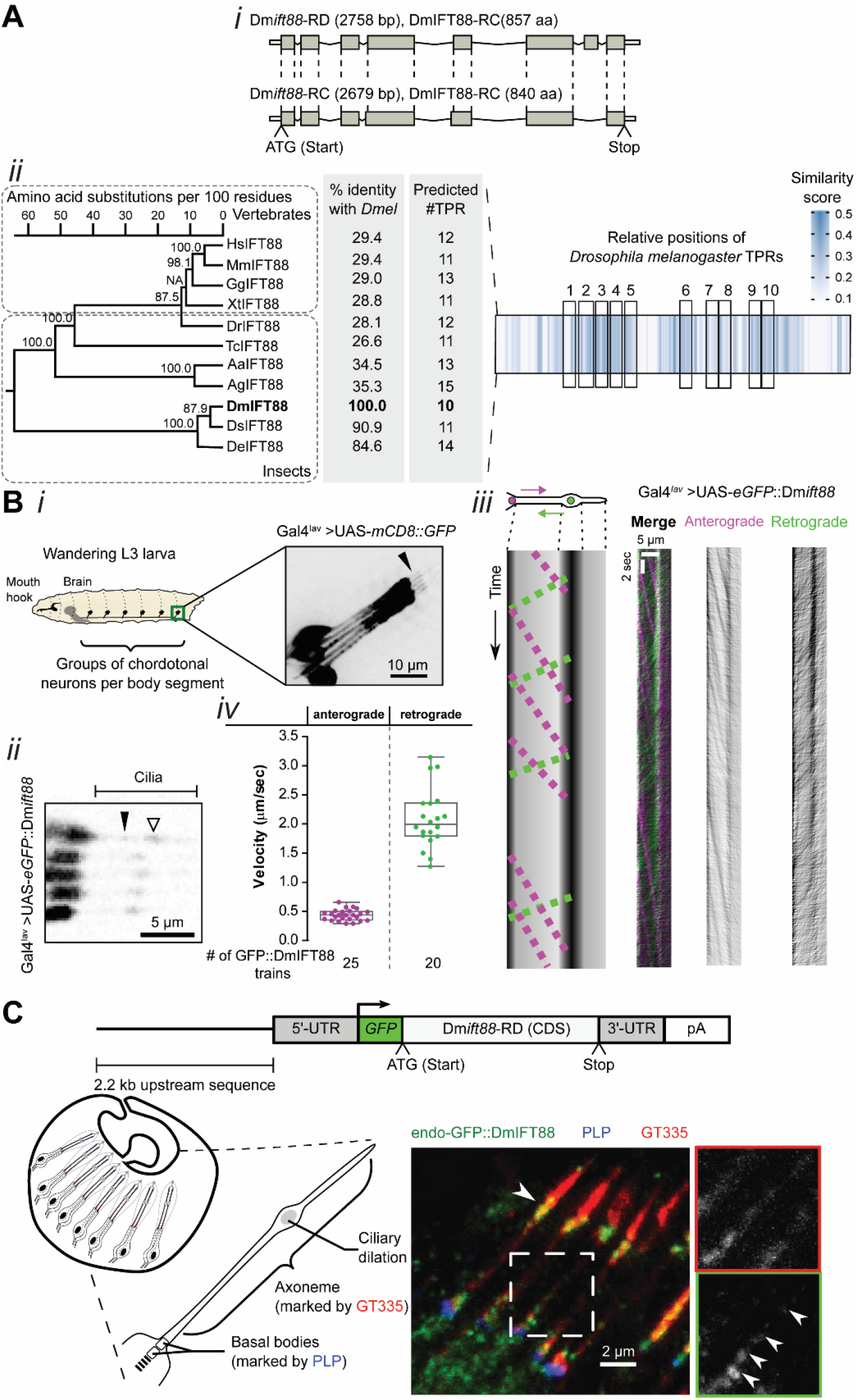
DmIFT88 is evolutionarily conserved and its trains are visible in *Drosophila* sensory cilia. **(A)** *Drosophila* IFT88 shows structural-domain conservation despite low amino acid sequence conservation. (i) Schematic representation of the two annotated Dm*IFT88* isoforms (RNA and proteins) in the fly. The grey boxes represent coding sequences. (ii) Left: Maximum-likelihood phylogenetic tree for IFT88 from various vertebrate and insect species, displaying bootstrap branch “support values” in percentages (%). The accession numbers of the proteins used in this analysis and a list of abbreviations are provided in the Supplemental Table 02. NA: the “support value” could not be calculated due to the method used to generate the sequence alignment. The amino acid identity of each sequence compared to *Drosophila melanogaster* is shown in percentages (%), and the number of predicted tetratricopeptide repeat (TPRs) domains in each species is displayed. Right: Multiple sequence alignment of IFT88 from eleven species represented as a heat map generated using JProfileGrid2. Each position in the alignment is shown as a box, colour-coded according to the similarity score. The relative positions of the ten TPRs of *Drosophila melanogaster* are indicated by black boxes. **(B)** GFP::DmIFT88 trains are visible in wandering L3 larvae. (i) Schematic representation of a L3 larva showing the segmentally arranged groups of chordotonal neurons (a group of five neurons is called lch5). Membrane-bound GFP (UAS-mCD8::GFP) is expressed using Gal4^*Iav*^ to visualise the morphology, including cilia (arrowhead), of one such group (lch5) of neurons. (ii) A movie still showing ectopically expressed GFP::DmIFT88 in the dendrite tip and cilia in lch5 neurons using Gal4^*Iav*^. Empty and filled arrowheads mark ciliary dilation and IFT particles, respectively. (iii) Left: A scheme of a cilium showing the IFT particles moving in anterograde (magenta) and retrograde (green) directions. Below the scheme, a hypothetical kymograph is shown that would result from the two types of particles moving in time along the cilium. Right: An example kymograph with DmIFT88 particles (colour-coded depending on their direction). The train-tracks were extracted using the *KymographClear* macro toolset from ImageJ. Vertical lines in all kymographs suggest that DmIFT88 also accumulates at the ciliary dilation. (iv) Quantifications of the speed of the anterograde and retrograde DmIFT88 particles, extracted from 20-25 movies from five different larvae. **(C)** DmIFT88 protein is found in fully formed cilia in the adult. Top: Representation of the transgene (from (Han et al., 2003)), expressing GFP::DmIFT88 near endogenous level, used to observe the DmIFT88 localisation in fly cilia. Left: A scheme of the chordotonal neuron architecture in the 2^nd^ antennal segment of adult flies showing the expected localisation of *Drosophila* Pericentrin-like protein (PLP) and glutamylated tubulin (GT335) in the basal bodies and the axoneme, respectively. Right: representative image of the localisation of endogenous GFP::DmIFT88 with respect to the two aforementioned markers in the adult chordotonal cilia. In C, GFP::DmIFT88 signals were enhanced with an anti-GFP antibody.

### DmIFT88 trains are visible in fully assembled cilia

To assess the localisation of DmIFT88 protein in fully assembled cilia, we first tested a transgenic fly line expressing GFP-tagged DmIFT88 protein (GFP::DmIFT88) under its endogenous promotor (Han et al., 2003), however its weak signal prevented its use for live imaging. We thus generated transgenic lines carrying a UAS-eGFP::DmIFT88 (isoform-RD) construct, as this isoform suffices to rescue IFT88 function in Dm*ift88/nompB* mutant in which all isoforms are affected (Han et al., 2003). To test whether DmIFT88 is part of IFT trains in assembled cilia, the transgene was expressed using a chordotonal neuron-specific driver (Gal4^*Iav*^) (Figure 1Bi) (Zhang et al., 2013). We were able to track DmIFT88 particles in the cilia of lateral chordotonal organ (lch5) neurons in wandering L3 larvae. Quantifications of the velocity of the eGFP::DmIFT88 signal (Figure 1Bii-iv and Supplemental movie 1) revealed that the particles move about five times faster in the retrograde direction (2.09 μm/s) than in the anterograde direction (0.44 μm/s). A recent study on the chordotonal neurons in developing *Drosophila* pupa found a similar anterograde velocity for IFT88 (∼0.44 µm/s), whereas retrograde velocities varied (∼0.12 and ∼0.7 µm/s) between ciliary compartments (Lee et al., 2018). These observations suggest that anterograde DmIFT88 velocities in different *Drosophila* chordotonal cilia types are largely constant, while retrograde velocities vary between developmental stages, cell types, and ciliary compartments. In fact, the IFT velocities are known to vary considerably between species and cell types of a given organism ((Besschetnova et al., 2010; Williams et al., 2014); an overview of different IFT velocities is provided in Supplemental Table 04). We also noticed that the signal intensities of the anterograde trains of eGFP::DmIFT88 appear stronger than the retrograde train intensities (Figure 1Biii). This result suggests that anterograde trains are longer than retrograde trains, similar to observations in other species (Fort et al., 2016; Pigino et al., 2009).

We next asked whether DmIFT88 is also present in chordotonal cilia of the 2^nd^ antennal segment in adults (Supplemental Figure 1B). We could detect both Dm*ift88* isoforms in mRNA isolated from adult antenna (Supplemental Figure 2B), suggesting this gene continues to be expressed after ciliogenesis. Because live imaging was not possible, we immuno-stained antennal sections of flies expressing GFP::DmIFT88 under endogenous promoter, and observed an enrichment of the protein at the ciliary base, as well as at the ciliary dilation of the chordotonal neurons (Figure 1C). The strong signals near the ciliary base and the dilation may represent an immobile fraction of the DmIFT88 pool, which is also supported by kymograph analysis (Figure 1Biii). Additional smaller and weaker GFP dots, similar to the ones observed in the larvae (Figure 1B, filled arrowhead), were detected along the axoneme, likely representing IFT trains (inset on Figure 1C). Together, our observations on both larval and adult cilia (Figure 1B, C) suggest that DmIFT88 is actively transported into the cilia beyond biogenesis.

**Figure 2:**
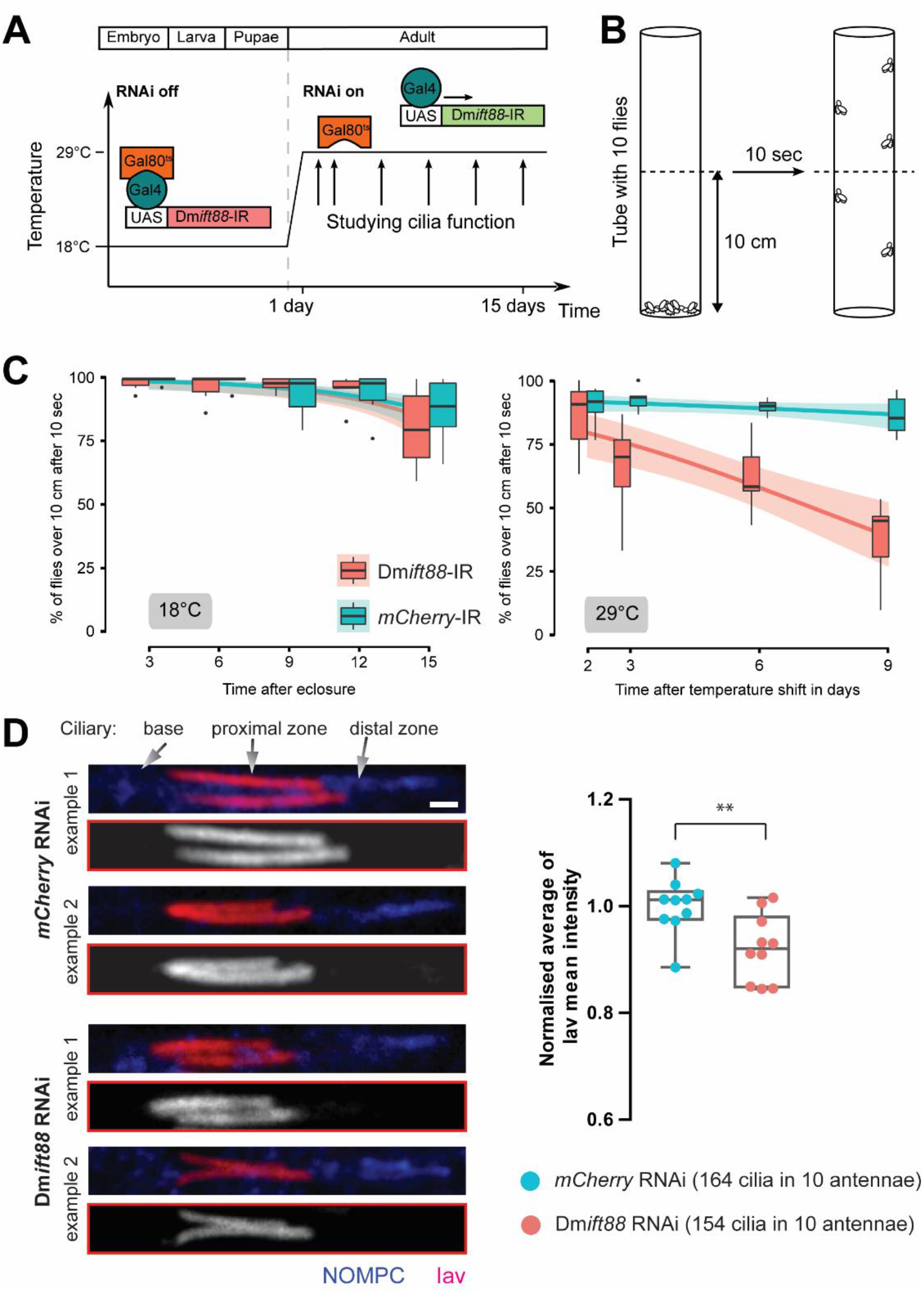
Acute removal of DmIFT88 in adult flies leads to impaired sensory functions and defective localisation of ciliary proteins. **(A)** Scheme of the approach and timeline of the conditional knockdown (Dm*ift88* and *mCherry* RNAis) experiments. Dm*ift88* and *mCherry* genes are knocked-down in cholinergic neurons, including chordotonal neurons, using Gal4^*Chat19b*^. Flies are reared at 18°C to repress the expression of the hairpin during development through the co-expression of a temperature-sensitive version of Gal80 ubiquitously (*Tub*Gal80^ts^). Upon eclosure, adult flies are shifted to 29°C (non-permissive temperature for Gal80^ts^) to activate the expression of RNA hairpins. **(B)** Schematic representation of the climbing (negative-gravitaxis) assay. The effect on sensory cilia function is approximated by quantifying climbing behaviour of the adult flies on specific days (arrows) after temperature shift. **(C)** Time-dependent changes in climbing behaviour at 18°C (left) and 29°C (right) in (control (*mCherry*) and Dm*ift88*) RNAi flies. Each boxplot corresponds to a total of 60 flies measured in sets of ten animals each. The data are fitted using linear regression. The curve fitted to the data is shown as a solid line, and the area around the curve represents the 95% confidence interval. The two lines are significantly different at 29°C, but no significant difference can be detected at 18°C. (**D**) Left: Representative immunofluorescence images of adult chordotonal cilia (with Inactive (Iav) and NOMPC in the proximal and distal zone of the cilia, respectively) from flies with control (*mCherry*) and Dm*ift88* RNAi. Scale bar: 1 µm. Right: Box plots of the normalised average Iav signal along the proximal part of the axoneme in flies with different genotypes. p-value is calculated using Mann-Whitney test (**-p-value < 0.01).

### Conditional knockdown of DmIFT88 in adult cilia causes climbing defects

We thus investigated whether DmIFT88 has any significant role in the maintenance of cilia. Since this protein is essential for ciliogenesis in sensory neurons (Han et al., 2003), we could not take advantage of the available mutant, as it lacks cilia. We thus used a Gal4-RNAi system to knockdown Dm*ift88* expression in a certain subset of sensory neurons only after cilia are fully formed. We first tested for the specificity and efficacy of the RNAi line (with a hairpin that targets an exon common to both Dm*ift88* isoforms) using a constitutively active promoter (Gal4^*Tub*^). We found that the mRNA levels of both isoforms are strongly reduced in antenna of flies expressing the Dm*ift88* hairpin (UAS-Dm*ift88*-IR), as compared to negative controls (UAS-*mCherry*-IR, as *mCherry* is not encoded in the fly genome; Supplemental Figure 2A and B). Importantly, sound stimulation evoked much weaker compound action potentials of the sound-sensitive antennal chordotonal neurons in DmIFT88 knockdown flies than in control flies (Supplemental Figure 2C), consistent with previously reported *Dmift88/nompB* mutant phenotypes (Han et al., 2003). Interestingly, we found that while flies with DmIFT88 knockdown do not assemble cilia, they can still build the transition zone (Supplemental Figure 2D-G), similar to the phenotype observed in *ift88* and *kinesin-2* mutants in various model organisms (Pazour et al., 2002; Sarpal et al., 2003).

As mentioned above, to test for defects in cilium maintenance, we used an inducible promotor system (Gal4^*Chat19b*^-*Tub*Gal80^ts^; explained in Figure 2A), which responds to the shift in temperature and can repress (18°) or allow for hairpin expression (29°) ((McGuire et al., 2003); for further details, see “Material and Methods” and Figure 2A). The Gal4^*Chat19b*^ driver expresses in the peripheral and central nervous system in all developmental stages (Salvaterra and Kitamoto, 2001). Since cilia biogenesis occurs during *Drosophila* development (Han et al., 2003), flies were first grown at 18°C, and only shifted to 29°C after eclosure, when ciliogenesis is complete. Because gravity sensing in *Drosophila* is mediated by chordotonal neurons, flies were then submitted to a behavioural (climbing) assay at different times for up to two weeks (Figure 2A, B; (Kamikouchi et al., 2009)). At 18°C, the flies with both experimental and control genotypes were nearly indistinguishable in their climbing performance (Figure 2C right), documenting that neither the genetic background nor the hairpins have behavioural adverse effects. Importantly, Dm*ift88* RNAi flies developed climbing defects within 3 days after shifting to 29°C (Figure 2C left). DmIFT88 is thus important for maintaining sensory behaviour after ciliogenesis is complete. Sensory defects caused by conditional DmIFT88 knockdown do not associate with morphological gross defects of chordotonal neuron cilia, yet, after 9 days post knockdown, the curvature of the base of cilia increased (Supplemental Figure 3). Interestingly, in larval *Drosophila* chordotonal neurons, the bending of the cilium at its base was described to be associated with cilium dysfunction (Zanini et al., 2018). Altogether, our data suggests that DmIFT88 is required for chordotonal cilia structure and function maintenance.

**Figure 3:**
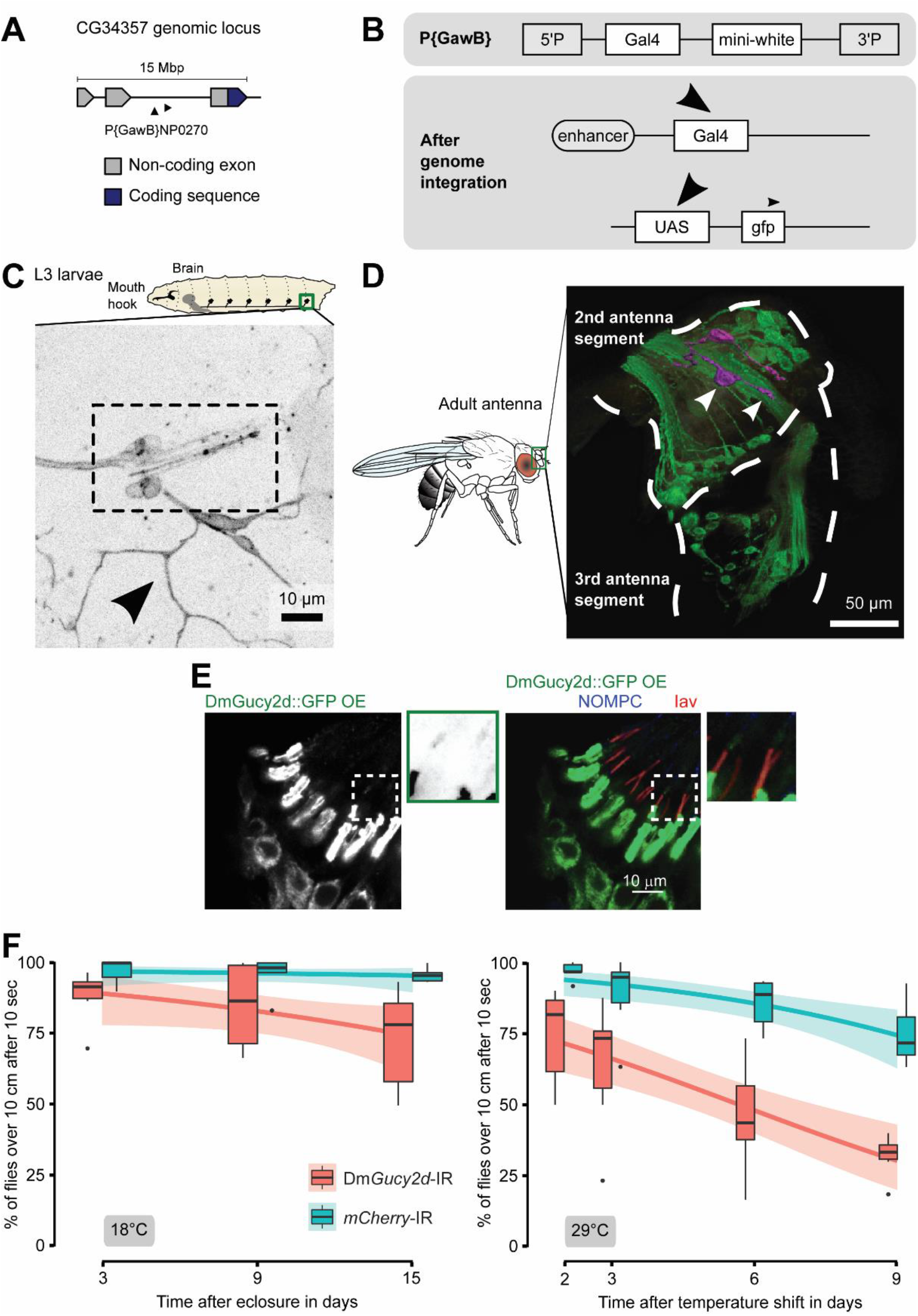
CG34357, a *Drosophila* homologue of Gucy2d, localises to chordotonal cilia and its removal impairs sensory function in adult flies. (**A**) Schematic presentation of the enhancer line used to determine *CG34357* expression. The P-element contains a Gal4 gene which was randomly inserted after *CG34357* exon 2. The upper box contains a schematic representation of the P{GawB} element showing two P-element arms flanking a Gal4 gene and a mini-white gene. The lower box shows schematically how it is used to determine *CG34357* expression pattern: presumably the same enhancers that activate *CG34357* also activate the Gal4 gene, which then drives expression of the GFP reporter gene inserted at the 3’-end of UAS. Using the enhancer line to drive UAS-mCD8::GFP expression, *CG34357* expression is detected in chordotonal neurons in L3 larvae (dashed box) as well as in other neurons, including non-ciliated class-I dendritic arborisation neurons, in the peripheral nervous system (arrowhead). (**D**) Fluorescence image shows *CG34357* expression pattern in the antennae of adult flies. The image stack was converted into a 3D model using Imaris software (see Supplemental video 02). The outline of the 2^nd^ and 3^rd^ antenna segment is drawn using the autofluorescence of the cuticle. Two chordotonal neurons in the second antenna segment are highlighted in magenta indicating the cell body (arrow) and the dendrite (arrowhead). **(E)** Immunofluorescence image shows that though the ectopically expressed DmGucy2d::GFP protein (using Gal4^*Iav*^) primarily accumulates in the dendrites of the adult chordotonal neurons, it also localises to the cilia. Insets (2x of the dashed boxes) show that the GFP signal can also be detected in the ciliary dilation (distal to the Iav protein signal). **(F)** Quantification of climbing behaviour in flies with conditional knockdown of Dm*Gucy2d* knockdown and *mCherry* (as internal control) at 18°C (left) and 29°C (right). Each boxplot corresponds to a total of 60 flies measured in sets of ten animals each. The data are fitted using linear regression. The curve fitted to the data is shown as a solid line, and the area around the curve represents the 95 % confidence interval. The two lines are significantly different at 29°C (experiment), but marginal difference can be detected at 18°C (control).

To assess whether the defects caused by DmIFT88 knockdown might be associated with an impairment in the transport of some components of the sensory apparatus, we stained the cilia with an antibody against the TRPV channel subunit, Inactive (Iav), which resides in the proximal cilium region of chordotonal neurons and is required for sound and gravity sensing (Sun et al., 2009). After 9 days of DmIFT88 knockdown, staining intensities were slightly, but significantly, reduced (Figure 2D). We further hypothesised that the climbing phenotype might also reflect mislocalisation of other unknown cargoes of DmIFT88, which are involved in signal transduction, besides Iav. Given that heterozygous mutations in *ift88* give raise to retinal degeneration in humans, it is possible that some of its cargoes would show a similar phenotype (Chekuri et al., 2018).

Interestingly, in IMCD3 cells, IFT88 transports ectopically expressed mouse Gucy2e, a membrane bound (particulate) cGMP-generating enzyme, to its primary cilium (Bhowmick et al., 2009, Besharse et al. 1990). Gucy2e is also essential for the function of vertebrate photoreceptors and is implicated in retina-specific ciliopathies that can be related to the development/maintenance of ciliary function, such as Leber congenital amaurosis (LCA) and retinitis pigmentosa (Zagel and Koch, 2014). Gucy2e seemed an interesting candidate DmIFT88 cargo as it is evolutionarily conserved, with homologues being found in several animals, such as *C. elegans*, mouse, and humans, where those signalling molecules regulate a wide range of sensory functions (reviewed in (Johnson and Leroux, 2010; Maruyama, 2016; Wen et al., 2014); Supplemental Table 03). We thus hypothesised that IFT88 might be generally involved in cilium maintenance in sensory neurons, in part by transporting particulate guanylyl cyclases such as Gucy2e.

### CG34357, a homologue of mouse Gucy2e, localises to chordotonal cilia and is required for maintaining their function in adult flies

We decided to search first for a *Drosophila melanogaster* homologue of Gucy2d/e and then to investigate its possible relation with DmIFT88. The *Drosophila* genome encodes several particulate guanylyl cyclases (Morton, 2004), but none of them has been previously associated with climbing behaviour, mechanosensation, or cilium function. Using the protein sequence of the mouse Gucy2e (an orthologue of human Gucy2d) in PSIBlastp search (Altschul et al., 1997) against the *Drosophila melanogaster* protein database, we identified several putative particulate guanylyl cyclases (Supplemental Figure 4A). We reasoned that knockdown of any guanylyl cyclase that functions in chordotonal sensory neurons would impair climbing behaviour. Given that RNAi reagents for four of those putative guanylyl cyclases were available, we used them to knockdown the respective genes using Gal4^*Chat19b*^. The climbing assays performed showed that the knockdown of three out of four candidates caused climbing defects, one of them being the gene *CG34357* (Supplemental Figure 4A). For follow-up studies, we focused on this gene because it is the closest *Drosophila* homologue of mouse *Gucy2e* and, accordingly, was named as Dm*Gucy2d*. To test if Dm*Gucy2d* is normally expressed in chordotonal neurons, we used an enhancer (Gal4)-trap line (NP0270), in which yeast Gal4 is inserted after the second exon 2 of Dm*Gucy2d* (Hayashi et al., 2002), to express a membrane-bound GFP reporter (UAS-mCD8::GFP) (Figure 3A and B). We observed that the enhancer is expressed in the chordotonal neurons in both L3 larvae and adult 2^nd^ antennal segment (Figure 3C and D). Notably, in the adult antenna, we detected GFP signal only in a few chordotonal neurons, suggesting that only a subset of chordotonal neurons express Dm*Gucy2d* or that the enhancer trap line might not fully recapitulate the endogenous gene expression (Figure 3D).

**Figure 4:**
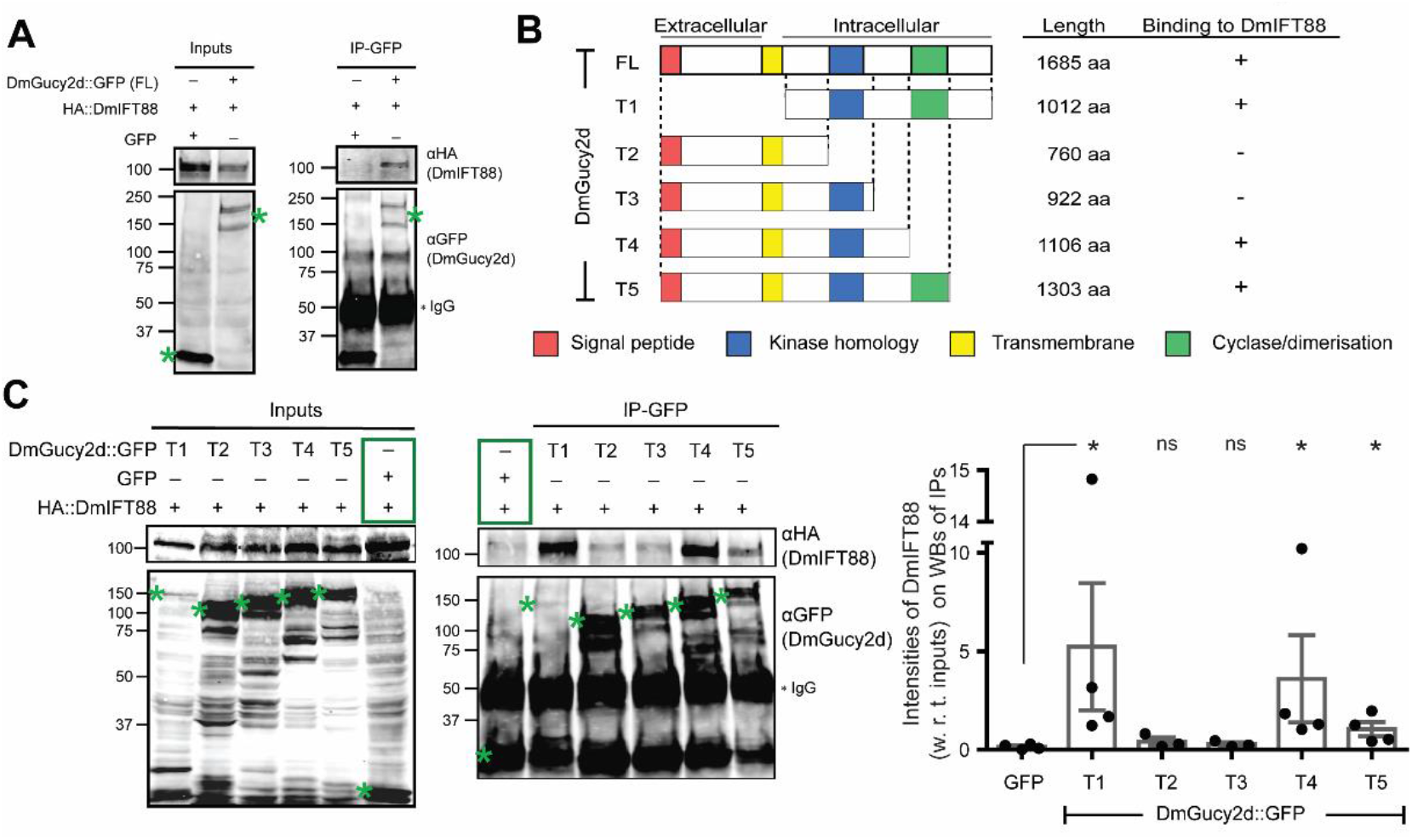
DmGucy2d binds DmIFT88 through its intracellular portion in cultured *Drosophila* (Dmel) cells. (**A**) Immunoprecipitation (IP) assay performed upon co-overexpression of 3xHA::DmIFT88, and DmGucy2d::GFP or GFP in Dmel cells. DmIFT88 co-immunoprecipitates with DmGucy2d::GFP, but not GFP alone. (**B**) Left: Schematic representation of DmGucy2d truncation constructs used to determine its binding region to DmIFT88, showing also the domain structures and constructs length (aa). Right: A summary of the various DmGucy2d truncated peptides ability to bind DmIFT88 (from C) is shown. (**C**) IP assay performed upon co-overexpression of full-length 3xHA::DmIFT88 and GFP-tagged fragments of DmGucy2d in Dmel cells. Right: Bar plots (overlaid with scattered dots) of the fraction of DmIFT88 (with respect to the inputs) binding to the GFP-tagged fragments of DmGucy2d and GFP alone. Each bar contains fractions of bound DmIFT88 intensity values (= values in the IP/ values in the Input) measured on the western blots, for at least three independent experiments. In A and C, co-overexpressed GFP and 3xHA::DmIFT88 serves as a negative control in the experiments (green boxes). Note that we also detect a faint non-specific HA-positive band in the IPs from extracts that co-express GFP and 3xHA::DmIFT88. The bands of expected molecular weight are marked with green asterisks. The p-values are calculated using Mann-Whitney tests (*-p-value < 0.05; ns-not significant).

The predicted protein product of DmGucy2d has all the features of a typical particulate guanylyl cyclase (i.e. signal peptide, transmembrane domain, kinase homology domain, dimerization domain and cyclase domain, see Supplemental Figure 4B and C). Three transcripts are expressed from the Dm*Gucy2d* gene locus and two of those differ only in the length of the 3’-untranslated regions (Dm*Gucy2d*-RD and -RC). We focused on the two longer transcripts as the third transcript leads to a small truncated protein that only contains the extracellular part of the protein and in principle should not be able to bind to DmIFT88, which is an intracellular protein. To further assess DmGucy2d localisation, we cloned the coding sequence of its longer isoform (RD/RC, as they only differ in their UTRs) and generated transgenic flies in which the construct was expressed under an UAS promotor. The protein was tagged with GFP at the C-terminus to avoid cleavage of the tag due to the existing signal peptide (Supplemental Figure 4B). When UAS-DmGucy2d::GFP was expressed using a chordotonal neuron-specific driver (Gal4^*Iav*^), GFP fluorescence was observed in the dendrites as well as along their cilia (Figure 3E).

We next wanted to test whether DmGucy2d is important for maintaining cilium function. We used the inducible promotor system (Gal4^*Chat19b*^-*Tub*Gal80^ts^) to express the hairpin against Dm*Gucy2d* in adult ciliated cells. The flies with UAS-Dm*Gucy2d* RNAi showed marginal reduction in their climbing performance already at 18°C (Figure 3F, right graph), possibly reflecting leaky expression of the construct. Importantly, after temperature shift to 29°C, flies with DmGucy2d knockdown developed climbing defects within 2 days, which became more severe after 9 days (Figure 3F, left graph). Collectively, our results suggest that DmGucy2d localises to adult chordotonal cilia and is needed for the maintenance of climbing behaviour.

### DmIFT88 binds to DmGucy2d intracellular domain and is required for the cyclase’ s ciliary localisation

To test whether DmGucy2d is a DmIFT88 cargo, we first examined whether DmGucy2d binds to DmIFT88. We used *Drosophila melanogaster* cultured (Dmel) cells, which normally express neither of these proteins nor any other IFT proteins, making them an appropriate system to test for the interactions of these proteins (Hu et al., 2017). We co-expressed either GFP-tagged DmGucy2d or GFP alone, together with HA-tagged DmIFT88. Unlike the GFP control, DmGucy2d::GFP (full length, FL) was able to bind to HA::DmIFT88 (Figure 4A). To narrow down the region of DmGucy2d required to bind DmIFT88, we generated several fragments of the cyclase. The T1 fragment contains the entire intracellular part of the protein. The other four fragments (T2-T5) contain the extracellular and the transmembrane domains (which might be necessary for the cyclases’s membrane localisation), as well as parts of the intracellular domain. T2-to-T5 fragments are successively longer with larger parts of the intracellular domain (Figure 4B). Our co-immunoprecipitation experiments revealed that T1, T4 and T5 co-immunoprecipitate DmIFT88 (Figure 4C), suggesting that a large part of the intracellular domain is required for the interaction.

Given that the T1 fragment of DmGucy2d encompasses the whole intracellular domain that interacts with DmIFT88, we next investigated this fragment’s localisation to the chordotonal neurons (Figure 4C). UAS-T1-DmGucy2d::GFP, when expressed in chordotonal neurons of adult flies using Gal4^*Iav*^, localised, like the full protein, to the dendrites and cilia of adult chordotonal cilia (Figure 3E and 5A). These data suggest that DmGucy2d intracellular domain plays a key role in its localisation, presumably through its transport by DmIFT88. To test whether DmIFT88 transports this cyclase to the cilia, we investigated the localization of GFP-tagged T1-fragment, while knocking down DmIFT88. We observed that the percentage of cilia with T1-DmGucy2d::GFP signal at the ciliary dilation, as well as the mean intensity of T1-DmGucy2d::GFP at the proximal axoneme, were significantly reduced in DmIFT88 knockdown flies when compared to controls, while the signal intensity at the cilia base remained unaffected (Figure 5B-F). These results show that DmIFT88 is required to transport the cyclase to the cilia.

**Figure 5:**
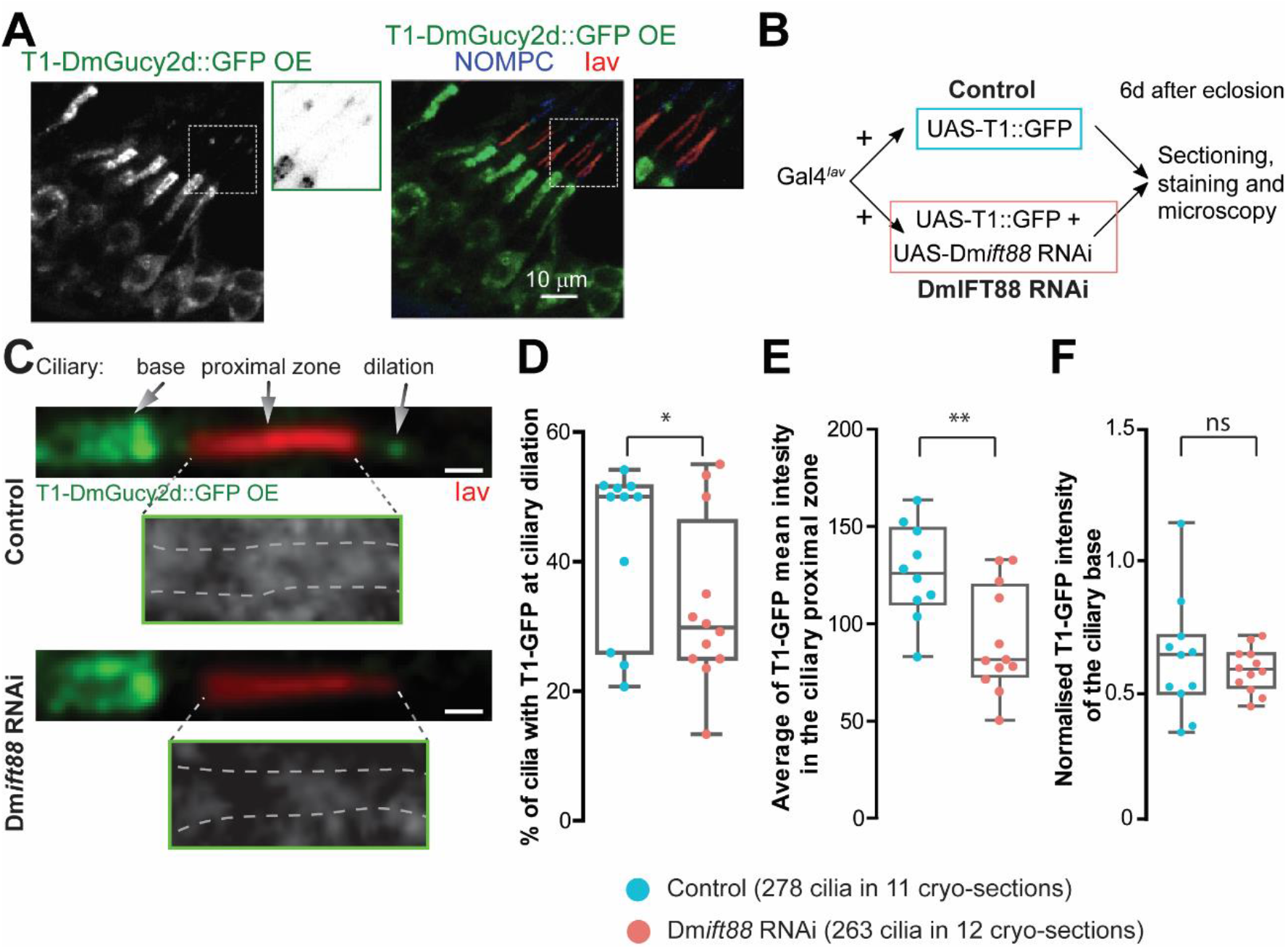
DmIFT88 is required for the ciliary localisation of DmGucy2d intracellular part. (**A**) Immunofluorescence images show that the ectopically expressed GFP-tagged T1-truncation of DmGucy2d (using Gal4^Iav^) accumulates in chordotonal neurons cell body, dendrite and ciliary dilation. Insets highlight that the GFP signal is also seen in the cilium. Proximal and distal cilia zones are marked with Iav and NOMPC antibodies, respectively. (**B**) Scheme summarises the experimental strategy in which T1-DmGucy2d::GFP is expressed in the chordotonal neurons (using Gal4^*Iav*^) with or without RNAi against Dm*ift88*. The resulting flies were analysed 6 days after eclosion. (**C**) Representative immunofluorescence images of the adult chordotonal cilia from flies with different genotypes (described in B). Insets show T1-GFP localisation along the proximal zone of the cilium (marked with dashed grey lines based on the anti-Iav antibody signal). Scale bars: 1 µm. (**D, E, F**) Box plots of the percentage of cilia with: GFP signal at the ciliary dilation (D), the average GFP signal along the proximal zone of the cilium (E), and normalised signal intensities of GFP at the ciliary base (F). The p-values are calculated using Welch corrected unpaired t-test (**-p-value < 0.01; *-p-value < 0.05; ns-not significant). Note that while no difference, in T1-DmGucy2d::GFP signal, is observed at the ciliary base, there is a clear difference both at the ciliary dilation and ciliary proximal zone.

In summary, we show that DmIFT88 binds the intracellular domain of DmGucy2d and transports it towards the ciliary dilation of the chordotonal cilia in adult flies. Both DmIFT88 and DmGucy2d are important for the maintenance of ciliary function. Given the importance of the intracellular domain of DmGucy2d for its transport we wondered whether this domain might be commonly mutated in disease. Indeed, in LCA patients, several mutations have been reported in the evolutionarily conserved intracellular domain of human Gucy2d (Supplemental Figure 5), suggesting that defective transport of Gucy2d into photoreceptor cilia might be one of the causes of the disease (de Castro-Miro et al., 2014; Feng et al., 2020; Jacobson et al., 2013; Li et al., 2011; Liu et al., 2020; Salehi Chaleshtori et al., 2020; Tucker et al., 2004; Zagel and Koch, 2014).

## Discussion and Conclusion

Cilia biogenesis depends on the activity of an exclusive transport system called IFT (Eguether et al., 2014; Jiang et al., 2015; Pazour et al., 2002), yet whether IFT also has a role in maintaining metazoan cilia had been little explored (Hao et al., 2011; Jiang et al., 2015; Marshall et al., 2005). Here we show that DmIFT88 expresses in- and is continuously mobile along the length of-fully formed sensory cilia. Furthermore, depletion of DmIFT88 after ciliogenesis in ciliated neurons leads to sensory behavioural phenotypes, suggesting a function for IFT in cilium maintenance, beyond its known role in ciliogenesis. We observe changes in the bending of the DmIFT88-depleted cilia at their bases, suggesting that IFT88 is required for maintaining the bending angle of the cilium. We also find a strong role for DmIFT88 in transporting Gucy2d, a particulate guanylyl cyclase, with a widespread signalling role in eukaryotes and an involvement in human disease. We additionally show that DmGucy2d localises to chordotonal cilia and its depletion in fully formed sensory neurons impairs behaviour (Figure 3). Finally, we demonstrate that DmIFT88 binds to the intracellular domain of DmGucy2d and is necessary for the cyclase’s ciliary localisation (Figure 4). Our findings identify a novel mechanism for ciliary maintenance.

### IFT88 plays differential roles in cilia assembly and maintenance

Multiple mechanisms have been implicated in the regulation of cilium homeostasis in diverse model organisms (Fort et al., 2016; Hao et al., 2011; Jiang et al., 2015; Marshall et al., 2005; Mourao et al., 2016). Ciliary structure and composition have to be maintained and they might alter in response to external stimuli, as found in *C. elegans* sensory cilia (DiTirro et al., 2019; Mykytyn and Askwith, 2017). Studies in *Chlamydomonas*, worm and mouse suggest that continuous tubulin turnover at the ciliary tip is required to maintain flagellum/cilium length (Hao et al., 2011; Jiang et al., 2015; Marshall et al., 2005). In contrast, in fruit flies, even though IFT88 is required for ciliary assembly (Han et al., 2003; Kohl et al., 2003; Pazour et al., 2002), we show that it has a mild role in maintaining the ciliary structure, while being critical for the maintenance of cilium function (Figure 1, 2). This result is similar to what was reported in *Trypanosoma*, suggesting that IFT88 plays an evolutionarily conserved role in the maintenance of the ciliary sensory function (Figure 2) (Fort et al., 2016).

Auditory chordotonal neurons are incredibly sensitive mechanical and sound receivers that can detect vibrations exerted by even Brownian motion (Nadrowski et al., 2008). DmGucy2d, shown here to be a ciliary cargo of DmIFT88, could contribute to signal amplification in multiple ways; on the one hand, DmGucy2d might be activated by low-level external stimulation, as Ca^2+^ regulates the activity of Gucy2d in mouse photoreceptor cells (Johnson and Leroux, 2010; Maruyama, 2016; Wen et al., 2014). On the other hand, it might also be involved in modifying the mechanical properties of the cilium in response to stimulation by adjusting the activities of dynein arms that are critical for mechanosensation of chordotonal cilia (Karak et al., 2015). Our data suggests that transport of DmGucy2d by DmIFT88 in fully formed sensory cilia is an important function of DmIFT88 in the maintenance of ciliary function. The mouse homologue of human Gucy2d, Gucy2e/GC-1, reportedly is transported by IFT88 into primary cilia in IMCD3 cells (Bhowmick et al., 2009), whereas this cyclase is transported into mouse photoreceptor cilia by rhodopsin (Pearring et al., 2015). It is thus possible that different modes of cyclase transport exist in different cell types and might even co-exist within the same cell. The fact that opsins localise to the dendrites of the fly chordotonal neurons and are required for their mechanotransduction function (Katana et al., 2019; Zanini et al., 2018) (Figure 1), makes it possible that opsins and DmIFT88 act together in DmGucy2d cilium localisation. To fully appreciate the potential of IFT in regulating various ciliary properties, such as structural maintenance and plasticity, as well as composition and function, it will be important to identify other cargoes that are transported by diverse IFT proteins in the future. Moreover, to fully understand how cilia are maintained, it will be important to uncover whether and how IFT-independent transport plays a role in that process.

### Implications of human Gucy2d evolutionarily conserved residues in its function and transport

Several mutations spread along the entire length of Gucy2d lead to Leber congenital amaurosis (LCA), a retina-specific ciliopathy (den Hollander et al., 2008). However, we know little about how those mutations cause the disease. Given that Gucy2d is a ciliary guanylyl cyclase, the mutations affecting either the protein’s stability, cyclase activity (Jacobson et al., 2013), or transport into the cilia can potentially be harmful. We uncovered that the intracellular domain of the cyclase, which we found to be essential for its transport into the cilium, is evolutionarily conserved. Furthermore, several of the Gucy2d residues, which are present in the intracellular part and mutated in patients, are conserved in *Drosophila* Gucy2d (Supplemental Figure 5 and Supplemental Table 8). These observations suggest that some of the phenotypes of LCA patients’ arise from insufficient delivery of Gucy2d to the photoreceptor cilia (de Castro-Miro et al., 2014; Feng et al., 2020; Jacobson et al., 2013; Li et al., 2011; Liu et al., 2020; Salehi Chaleshtori et al., 2020; Tucker et al., 2004; Zagel and Koch, 2014). Further studies to test this prediction would help to better understand Gucy2d transport and function, as well as the cellular basis of LCA and other related diseases.

## Acknowledgements

We thank Daniel F. Eberl, Li E. Cheng, Changsoo Kim and Elio Sucena for reagents. We thank Lab members for reviewing the manuscript and providing helpful discussions on the manuscript. We thank the IGC Advanced Imaging unit (and its Head, Gabriel G. Martins) and IGC Electron Microscopy unit (and its Head, Erin Tranfield) for helping us with image acquisition, and the IGC fly facility for assisting us with fly husbandry. SW, POR and SCJ are supported by the FCT (Fundação Portuguesa para a Ciência e Tecnologia) Fellowships/Grants/Contracts SFRH/BD/52176/2013, PTDC/BIA-BID/32225/2017 and SFRH/BPD/87479/2012, respectively. SCJ and MBD acknowledge FCT (PTDC/BIA-CEL/32631/2017 to SCJ) and European Research Council Consolidator Grant (CoG683528 to MBD) for their support through research grants.

## Notes

### Competing Interest Statement

The authors have declared no competing interest.

## References

Allan, R.K., and T. Ratajczak. 2011. Versatile TPR domains accommodate different modes of target protein recognition and function. Cell stress & chaperones. 16:353–367.

Altschul, S.F., T.L. Madden, A.A. Schaffer, J. Zhang, Z. Zhang, W. Miller, and D.J. Lipman. 1997. Gapped BLAST and PSI-BLAST: a new generation of protein database search programs. Nucleic acids research. 25:3389–3402.

Avidor-Reiss, T., A.M. Maer, E. Koundakjian, A. Polyanovsky, T. Keil, S. Subramaniam, and C.S. Zuker. 2004. Decoding cilia function: defining specialized genes required for compartmentalized cilia biogenesis. Cell. 117:527–539.

Besschetnova, T.Y., E. Kolpakova-Hart, Y. Guan, J. Zhou, B.R. Olsen, and J.V. Shah. 2010. Identification of signaling pathways regulating primary cilium length and flow-mediated adaptation. Current biology: CB. 20:182–187.

Bhowmick, R., M. Li, J. Sun, S.A. Baker, C. Insinna, and J.C. Besharse. 2009. Photoreceptor IFT complexes containing chaperones, guanylyl cyclase 1 and rhodopsin. Traffic. 10:648–663.

Breslow, D.K., and A.J. Holland. 2019. Mechanism and Regulation of Centriole and Cilium Biogenesis. Annual review of biochemistry. 88:691–724.

Chekuri, A., A.A. Guru, P. Biswas, K. Branham, S. Borooah, A. Soto-Hermida, M. Hicks, N.W. Khan, H. Matsui, A. Alapati, P.B. Raghavendra, S. Roosing, S. Sarangapani, S. Mathavan, A. Telenti, J.R. Heckenlively, S.A. Riazuddin, K.A. Frazer, P.A. Sieving, and R. Ayyagari. 2018. IFT88 mutations identified in individuals with non-syndromic recessive retinal degeneration result in abnormal ciliogenesis. Human genetics. 137:447–458.

Chintapalli, V.R., J. Wang, and J.A. Dow. 2007. Using FlyAtlas to identify better Drosophila melanogaster models of human disease. Nature genetics. 39:715–720.

de Castro-Miro, M., E. Pomares, L. Lores-Motta, R. Tonda, J. Dopazo, G. Marfany, and R. Gonzalez-Duarte. 2014. Combined genetic and high-throughput strategies for molecular diagnosis of inherited retinal dystrophies. PloS one. 9:e88410.

DiTirro, D., A. Philbrook, K. Rubino, and P. Sengupta. 2019. The Caenorhabditis elegans Tubby homolog dynamically modulates olfactory cilia membrane morphogenesis and phospholipid composition. eLife. 8.

Eguether, T., J.T. San Agustin, B.T. Keady, J.A. Jonassen, Y. Liang, R. Francis, K. Tobita, C.A. Johnson, Z.A. Abdelhamed, C.W. Lo, and G.J. Pazour. 2014. IFT27 links the BBSome to IFT for maintenance of the ciliary signaling compartment. Developmental cell. 31:279–290.

Fan, Z.C., R.H. Behal, S. Geimer, Z. Wang, S.M. Williamson, H. Zhang, D.G. Cole, and H. Qin. 2010. Chlamydomonas IFT70/CrDYF-1 is a core component of IFT particle complex B and is required for flagellar assembly. Molecular biology of the cell. 21:2696–2706.

Feng, X., T. Wei, J. Sun, Y. Luo, Y. Huo, P. Yu, J. Chen, X. Wei, M. Qi, and Y. Ye. 2020. The pathogenicity of novel GUCY2D mutations in Leber congenital amaurosis 1 assessed by HPLC-MS/MS. PloS one. 15:e0231115.

Fernández-Hernández, I., E.B. Marsh, and M.A. Bonaguidi. 2020. Mechanosensory neuron regeneration in adult < emϭ rosophila< /em>. bioRxiv:812057.

Fort, C., S. Bonnefoy, L. Kohl, and P. Bastin. 2016. Intraflagellar transport is required for the maintenance of the trypanosome flagellum composition but not its length. Journal of cell science. 129:3026–3041.

Grotewiel, M.S., I. Martin, P. Bhandari, and E. Cook-Wiens. 2005. Functional senescence in Drosophila melanogaster. Ageing research reviews. 4:372–397.

Han, Y.G., B.H. Kwok, and M.J. Kernan. 2003. Intraflagellar transport is required in Drosophila to differentiate sensory cilia but not sperm. Current biology: CB. 13:1679–1686.

Hao, L., M. Thein, I. Brust-Mascher, G. Civelekoglu-Scholey, Y. Lu, S. Acar, B. Prevo, S. Shaham, and J.M. Scholey. 2011. Intraflagellar transport delivers tubulin isotypes to sensory cilium middle and distal segments. Nature cell biology. 13:790–798.

Hayashi, S., K. Ito, Y. Sado, M. Taniguchi, A. Akimoto, H. Takeuchi, T. Aigaki, F. Matsuzaki, H. Nakagoshi, T. Tanimura, R. Ueda, T. Uemura, M. Yoshihara, and S. Goto. 2002. GETDB, a database compiling expression patterns and molecular locations of a collection of Gal4 enhancer traps. Genesis. 34:58–61.

Hu, Y., A. Comjean, N. Perrimon, and S.E. Mohr. 2017. The Drosophila Gene Expression Tool (DGET) for expression analyses. BMC bioinformatics. 18:98.

Jacobson, S.G., A.V. Cideciyan, I.V. Peshenko, A. Sumaroka, E.V. Olshevskaya, L. Cao, S.B. Schwartz, A.J. Roman, M.B. Olivares, S. Sadigh, K.W. Yau, E. Heon, E.M. Stone, and A.M. Dizhoor. 2013. Determining consequences of retinal membrane guanylyl cyclase (RetGC1) deficiency in human Leber congenital amaurosis en route to therapy: residual cone-photoreceptor vision correlates with biochemical properties of the mutants. Human molecular genetics. 22:168–183.

Jana, S.C., M. Girotra, and K. Ray. 2011. Heterotrimeric kinesin-II is necessary and sufficient to promote different stepwise assembly of morphologically distinct bipartite cilia in Drosophila antenna. Molecular biology of the cell. 22:769–781.

Jiang, L., Y. Wei, C.C. Ronquillo, R.E. Marc, B.K. Yoder, J.M. Frederick, and W. Baehr. 2015. Heterotrimeric kinesin-2 (KIF3) mediates transition zone and axoneme formation of mouse photoreceptors. The Journal of biological chemistry. 290:12765–12778.

Johnson, J.L., and M.R. Leroux. 2010. cAMP and cGMP signaling: sensory systems with prokaryotic roots adopted by eukaryotic cilia. Trends in cell biology. 20:435–444.

Kamikouchi, A., H.K. Inagaki, T. Effertz, O. Hendrich, A. Fiala, M.C. Gopfert, and K. Ito. 2009. The neural basis of Drosophila gravity-sensing and hearing. Nature. 458:165–171.

Karak, S., J.S. Jacobs, M. Kittelmann, C. Spalthoff, R. Katana, E. Sivan-Loukianova, M.A. Schon, M.J. Kernan, D.F. Eberl, and M.C. Gopfert. 2015. Diverse Roles of Axonemal Dyneins in Drosophila Auditory Neuron Function and Mechanical Amplification in Hearing. Scientific reports. 5:17085.

Karpenahalli, M.R., A.N. Lupas, and J. Soding. 2007. TPRpred: a tool for prediction of TPR-, PPR- and SEL1-like repeats from protein sequences. BMC bioinformatics. 8:2.

Katana, R., C. Guan, D. Zanini, M.E. Larsen, D. Giraldo, B.R.H. Geurten, C.F. Schmidt, S.G. Britt, and M.C. Gopfert. 2019. Chromophore-Independent Roles of Opsin Apoproteins in Drosophila Mechanoreceptors. Current biology: CB. 29:2961–2969 e2964.

Kohl, L., D. Robinson, and P. Bastin. 2003. Novel roles for the flagellum in cell morphogenesis and cytokinesis of trypanosomes. The EMBO journal. 22:5336–5346.

Lee, E., E. Sivan-Loukianova, D.F. Eberl, and M.J. Kernan. 2008. An IFT-A protein is required to delimit functionally distinct zones in mechanosensory cilia. Current biology: CB. 18:1899–1906.

Lee, N., J. Park, Y.C. Bae, J.H. Lee, C.H. Kim, and S.J. Moon. 2018. Time-Lapse Live-Cell Imaging Reveals Dual Function of Oseg4, Drosophila WDR35, in Ciliary Protein Trafficking. Molecules and cells. 41:676–683.

Li, A., M. Saito, J.Z. Chuang, Y.Y. Tseng, C. Dedesma, K. Tomizawa, T. Kaitsuka, and C.H. Sung. 2011. Ciliary transition zone activation of phosphorylated Tctex-1 controls ciliary resorption, S-phase entry and fate of neural progenitors. Nature cell biology. 13:402–411.

Liu, X., K. Fujinami, K. Kuniyoshi, M. Kondo, S. Ueno, T. Hayashi, K. Mochizuki, S. Kameya, L. Yang, Y. Fujinami-Yokokawa, G. Arno, N. Pontikos, H. Sakuramoto, T. Kominami, H. Terasaki, S. Katagiri, K. Mizobuchi, N. Nakamura, K. Yoshitake, Y. Miyake, S. Li, T. Kurihara, K. Tsubota, T. Iwata, K. Tsunoda, and C. Japan Eye Genetics. 2020. Clinical and Genetic Characteristics of 15 Affected Patients From 12 Japanese Families with GUCY2D-Associated Retinal Disorder. Translational vision science & technology. 9:2.

Marshall, W.F., H. Qin, M. Rodrigo Brenni, and J.L. Rosenbaum. 2005. Flagellar length control system: testing a simple model based on intraflagellar transport and turnover. Molecular biology of the cell. 16:270–278.

Maruyama, I.N. 2016. Receptor Guanylyl Cyclases in Sensory Processing. Frontiers in endocrinology. 7:173.

McGuire, S.E., P.T. Le, A.J. Osborn, K. Matsumoto, and R.L. Davis. 2003. Spatiotemporal rescue of memory dysfunction in Drosophila. Science. 302:1765–1768.

Milenkovic, L., M.P. Scott, and R. Rohatgi. 2009. Lateral transport of Smoothened from the plasma membrane to the membrane of the cilium. The Journal of cell biology. 187:365–374.

Morton, D.B. 2004. Atypical soluble guanylyl cyclases in Drosophila can function as molecular oxygen sensors. The Journal of biological chemistry. 279:50651–50653.

Mourao, A., S.T. Christensen, and E. Lorentzen. 2016. The intraflagellar transport machinery in ciliary signaling. Current opinion in structural biology. 41:98–108.

Mykytyn, K., and C. Askwith. 2017. G-Protein-Coupled Receptor Signaling in Cilia. Cold Spring Harbor perspectives in biology. 9.

Nadrowski, B., J.T. Albert, and M.C. Gopfert. 2008. Transducer-based force generation explains active process in Drosophila hearing. Current biology: CB. 18:1365–1372.

Pazour, G.J., S.A. Baker, J.A. Deane, D.G. Cole, B.L. Dickert, J.L. Rosenbaum, G.B. Witman, and J.C. Besharse. 2002. The intraflagellar transport protein, IFT88, is essential for vertebrate photoreceptor assembly and maintenance. The Journal of cell biology. 157:103–113.

Pearring, J.N., W.J. Spencer, E.C. Lieu, and V.Y. Arshavsky. 2015. Guanylate cyclase 1 relies on rhodopsin for intracellular stability and ciliary trafficking. eLife. 4.

Pigino, G., S. Geimer, S. Lanzavecchia, E. Paccagnini, F. Cantele, D.R. Diener, J.L. Rosenbaum, and P. Lupetti. 2009. Electron-tomographic analysis of intraflagellar transport particle trains in situ. The Journal of cell biology. 187:135–148.

Reiter, J.F., and M.R. Leroux. 2017. Genes and molecular pathways underpinning ciliopathies. Nature reviews. Molecular cell biology. 18:533–547.

Salehi Chaleshtori, A.R., M. Garshasbi, A. Salehi, and M. Noruzinia. 2020. The Identification and Stereochemistry Analysis of a Novel Mutation p.(D367Tfs*61) in the CYP1B1 Gene: A Case Report. Journal of current ophthalmology. 32:114–118.

Salvaterra, P.M., and T. Kitamoto. 2001. Drosophila cholinergic neurons and processes visualized with Gal4/UAS-GFP. Brain research. Gene expression patterns. 1:73–82.

Sarpal, R., S.V. Todi, E. Sivan-Loukianova, S. Shirolikar, N. Subramanian, E.C. Raff, J.W. Erickson, K. Ray, and D.F. Eberl. 2003. Drosophila KAP interacts with the kinesin II motor subunit KLP64D to assemble chordotonal sensory cilia, but not sperm tails. Current biology: CB. 13:1687–1696.

Sreekumar, V., and D.P. Norris. 2019. Cilia and development. Current opinion in genetics & development. 56:15–21.

Taschner, M., S. Bhogaraju, and E. Lorentzen. 2012. Architecture and function of IFT complex proteins in ciliogenesis. Differentiation; research in biological diversity. 83:S12–22.

Tucker, C.L., V. Ramamurthy, A.L. Pina, M. Loyer, S. Dharmaraj, Y. Li, I.H. Maumenee, J.B. Hurley, and R.K. Koenekoop. 2004. Functional analyses of mutant recessive GUCY2D alleles identified in Leber congenital amaurosis patients: protein domain comparisons and dominant negative effects. Molecular vision. 10:297–303.

van der Burght, S.N., S. Rademakers, J.L. Johnson, C. Li, G.J. Kremers, A.B. Houtsmuller, M.R. Leroux, and G. Jansen. 2020. Ciliary Tip Signaling Compartment Is Formed and Maintained by Intraflagellar Transport. Current biology: CB. 30:4299–4306 e4295.

Wen, X.H., A.M. Dizhoor, and C.L. Makino. 2014. Membrane guanylyl cyclase complexes shape the photoresponses of retinal rods and cones. Frontiers in molecular neuroscience. 7:45.

Williams, C.L., J.C. McIntyre, S.R. Norris, P.M. Jenkins, L. Zhang, Q. Pei, K. Verhey, and J.R. Martens. 2014. Direct evidence for BBSome-associated intraflagellar transport reveals distinct properties of native mammalian cilia. Nature communications. 5:5813.

Ye, F., D.K. Breslow, E.F. Koslover, A.J. Spakowitz, W.J. Nelson, and M.V. Nachury. 2013. Single molecule imaging reveals a major role for diffusion in the exploration of ciliary space by signaling receptors. eLife. 2:e00654.

Zagel, P., and K.W. Koch. 2014. Dysfunction of outer segment guanylate cyclase caused by retinal disease related mutations. Frontiers in molecular neuroscience. 7:4.

Zanini, D., D. Giraldo, B. Warren, R. Katana, M. Andres, S. Reddy, S. Pauls, N. Schwedhelm-Domeyer, B.R.H. Geurten, and M.C. Gopfert. 2018. Proprioceptive Opsin Functions in Drosophila Larval Locomotion. Neuron. 98:67–74 e64.

Zhang, W., Z. Yan, L.Y. Jan, and Y.N. Jan. 2013. Sound response mediated by the TRP channels NOMPC, NANCHUNG, and INACTIVE in chordotonal organs of Drosophila larvae. Proceedings of the National Academy of Sciences of the United States of America. 110:13612–13617.

