## Supplemental Information (Figures, Tables, Materials and Methods) for "IFT88 transports Gucy2d, a guanylyl cyclase, to maintain sensory cilia function in *Drosophila*"

### Supplemental Figures and Legends:

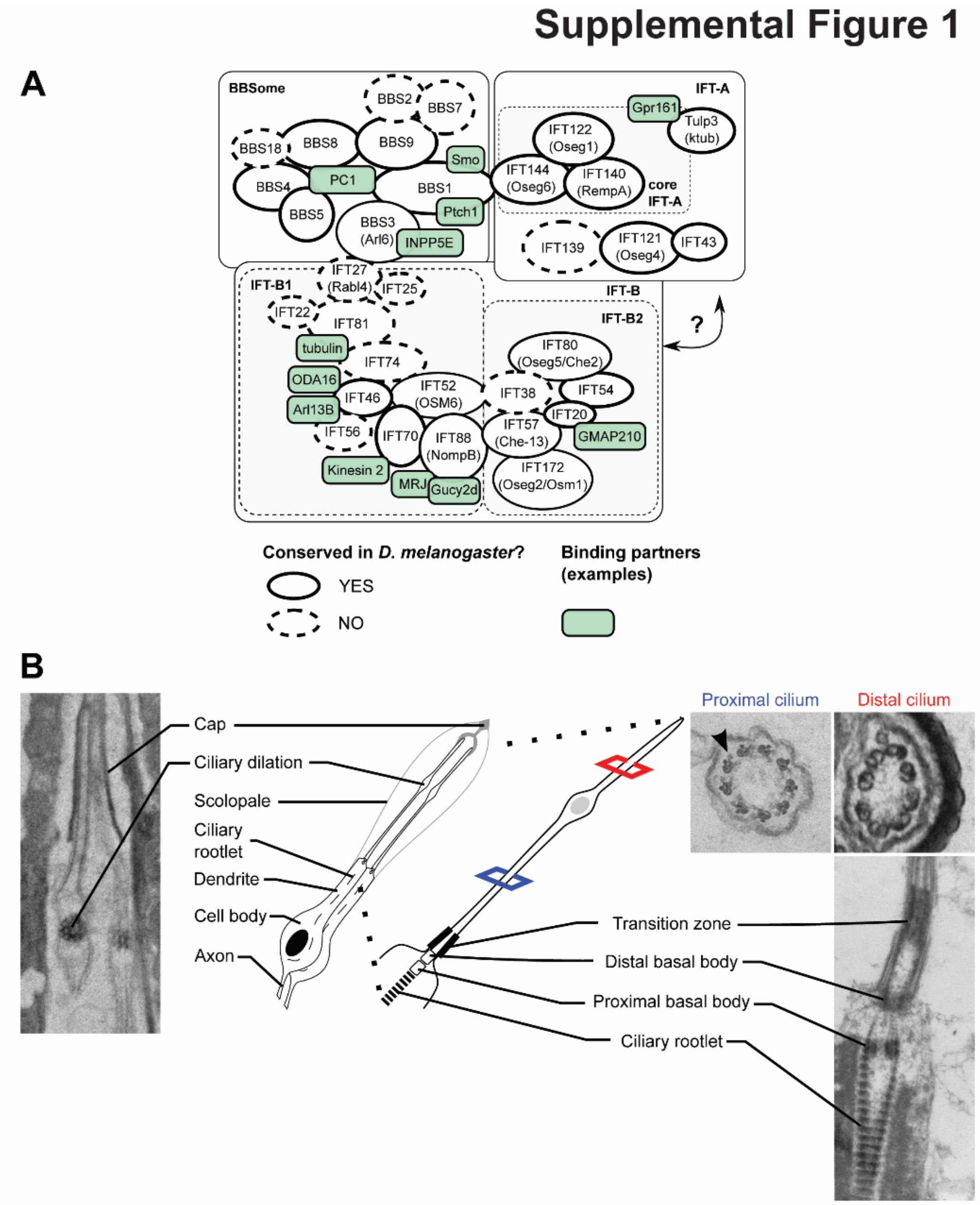


**Supplemental Figure 1: Conservation of the Intraflagellar Transport (IFT) complex in *Drosophila* and description of the ultrastructure of chordotonal cilia in the adult 2$^{nd}$ antennal segment. (A)** Schematic presentation of the components of the IFT complex found in human, *Chlamydomonas* (*C. reinhardtii*) and *Drosophila* (*D. melanogaster*). IFT consists of three complexes: IFT-A, IFT-B and the BBsome. IFT-A and –B are further divided based on their genetic features and biochemical properties. For the IFT-A and –B proteins, the *Chlamydomonas* nomenclature is used. Proteins conserved and not conserved in *Drosophila* are encircled by full and dashed outlines, respectively. The scheme also shows where the three complexes interact and highlights some known binding partners (green boxes). This scheme is modified from (Mourao et al., 2016). **(B)** Ultrastructural organisation of the cilium of chordotonal neurons in the fly 2nd antennal segment, as seen by electron micrographs. Here, two neurons project their cilia towards a cap cell. Each ciliary shaft is divided by the ciliary dilation into a proximal and a distal zone, which have distinct axonemal architectures, with only the proximal zone harbouring axonemal dynein arms (arrowhead). The base of the cilia is formed by distal and proximal basal bodies and the rootlet (Jana et al., 2016).

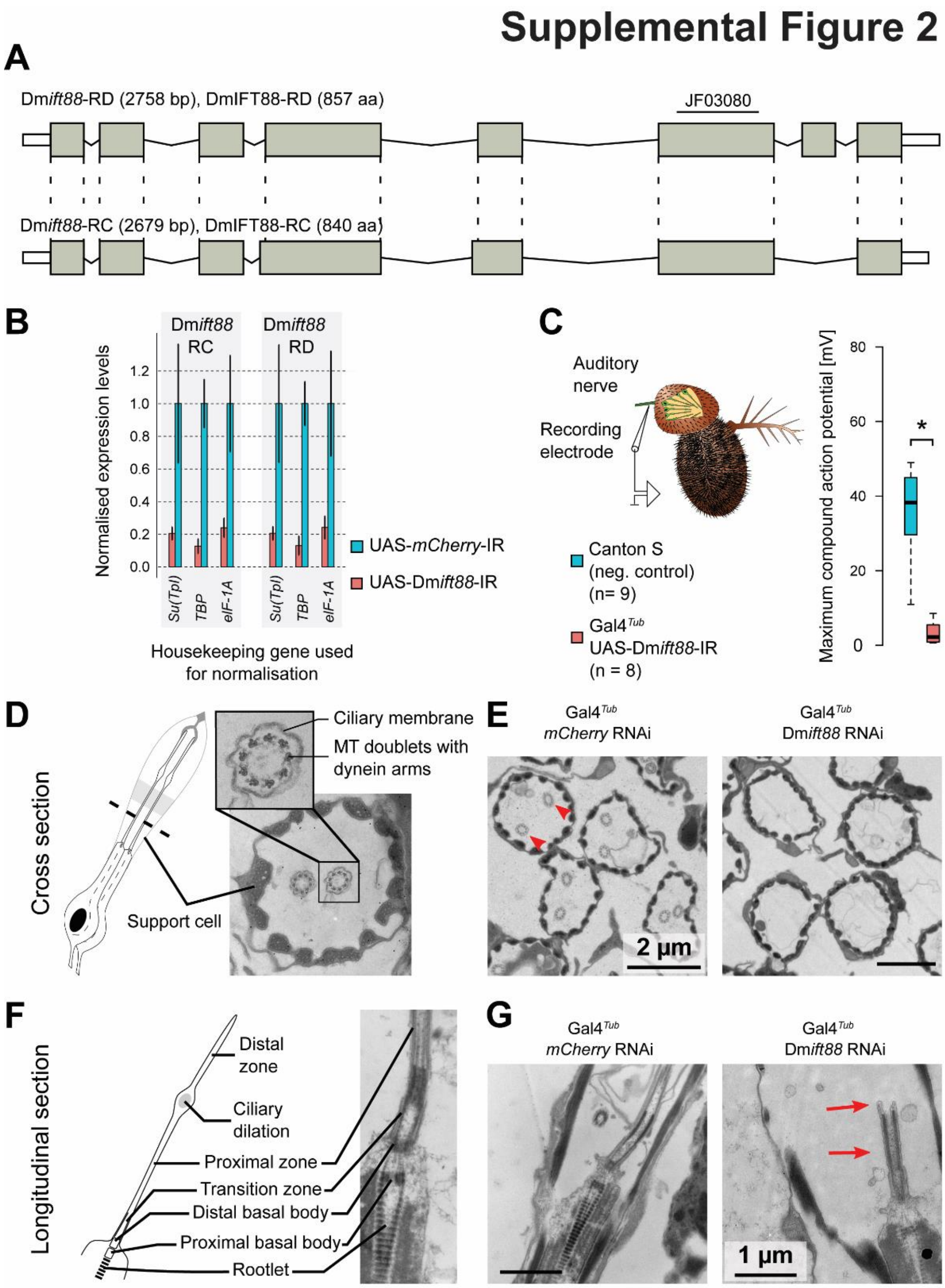


**Supplemental Figure 2: Validation of the Dm*ift88* inverted repeat (IR) used to downregulate Dm*ift88* expression. (A)** Schematic representation showing that the hairpin (ID: JF03080) affects both Dm*ift88* isoforms by targeting a common exon. **(B)** Bar plots (Mean ± S.D.) of real-time PCR quantification of Dm*ift88* isoform levels. mRNA was extracted from the antennae of flies expressing a hairpin against either *mCherry* (negative control) or Dm*ift88*. The hairpin expression was driven using the tubulin promoter (Gal4$^{Tub}$). The total mRNA was extracted three times per hairpin and each RNA sample was measured in triplicates. The data is normalised using three different housekeeping genes (*Su(Tpl)*, *TBP* and *eIF-1A*) to account for potential transcriptional changes of housekeeping genes as a consequence of the knockdown. **(C)** Box plots of maximum sound-evoked compound action potentials of chordotonal neurons in the second antennal segment of wild type flies (Canton S; negative control) and DmIFT88 knockdown flies expressing constitutively Dm*ift88*-IR ($Gal4^{Tub}$>UAS-Dm*ift88* RNAi). The sound responses are virtually abolished from the antenna of DmIFT88 knockdown flies (*p-value = 6.274e$^{-05}$ from Welch two sample t-test). Action potentials are measured from 4-6 days old flies. Median ± full range of variation (minimum to maximum). **(D-G)** The flies with knockdown on Dm*ift88* using ubiquitously promoter ($Gal4^{Tub}$) do not grow chordotonal cilia. Schematic representation and the representative EM images of wild type chordotonal cilia in cross (D) and longitudinal section (F). Representative images of cross-sections (E) and longitudinal-sections (G) of the chordotonal cilia in the adult flies ubiquitously ($Gal4^{Tub}$) expressing *mCherry*-IR (negative control) or Dm*ift88*-IR. Note that upon ubiquitous knockdown of Dm*ift88*, chordotonal neurons form ciliary transition zones (red arrows), but fail to extend cilia.

### Supplemental Figure 3

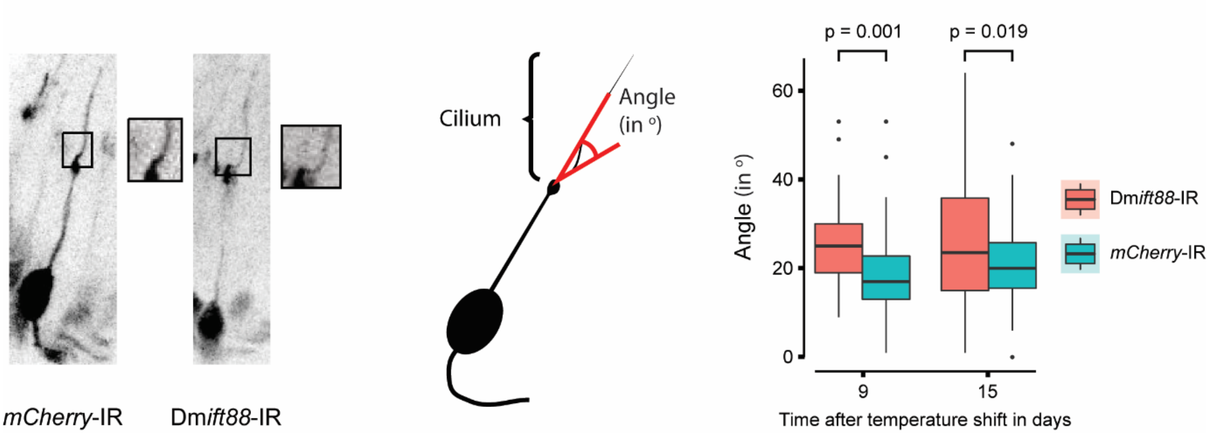


**Supplemental Figure 3: Continuous knockdown of DmIFT88 in the adult increases the curvature of the axoneme at the base of the cilium**. Left: Representative images of chordotonal neurons in the 2nd antennal segment of flies with mCherry or DmIFT88 knockdown for 15 days. Neurons are visualised by co-expressing GFP together with the respective hairpin. Long term knockdown of DmIFT88 affects ciliary (axoneme) bending (scheme in the Middle): cilia in DmIFT88 knockdown flies project out at a different angle from the dendrites than cilia in the control flies, as shown in the box plots (right). Data is obtained from antenna whole mounts to avoid artefacts from the sectioning process. Each box plot corresponds to the value of 50 angles from 5 different antennae. p-values are calculated using Welch Two Sample t-test.

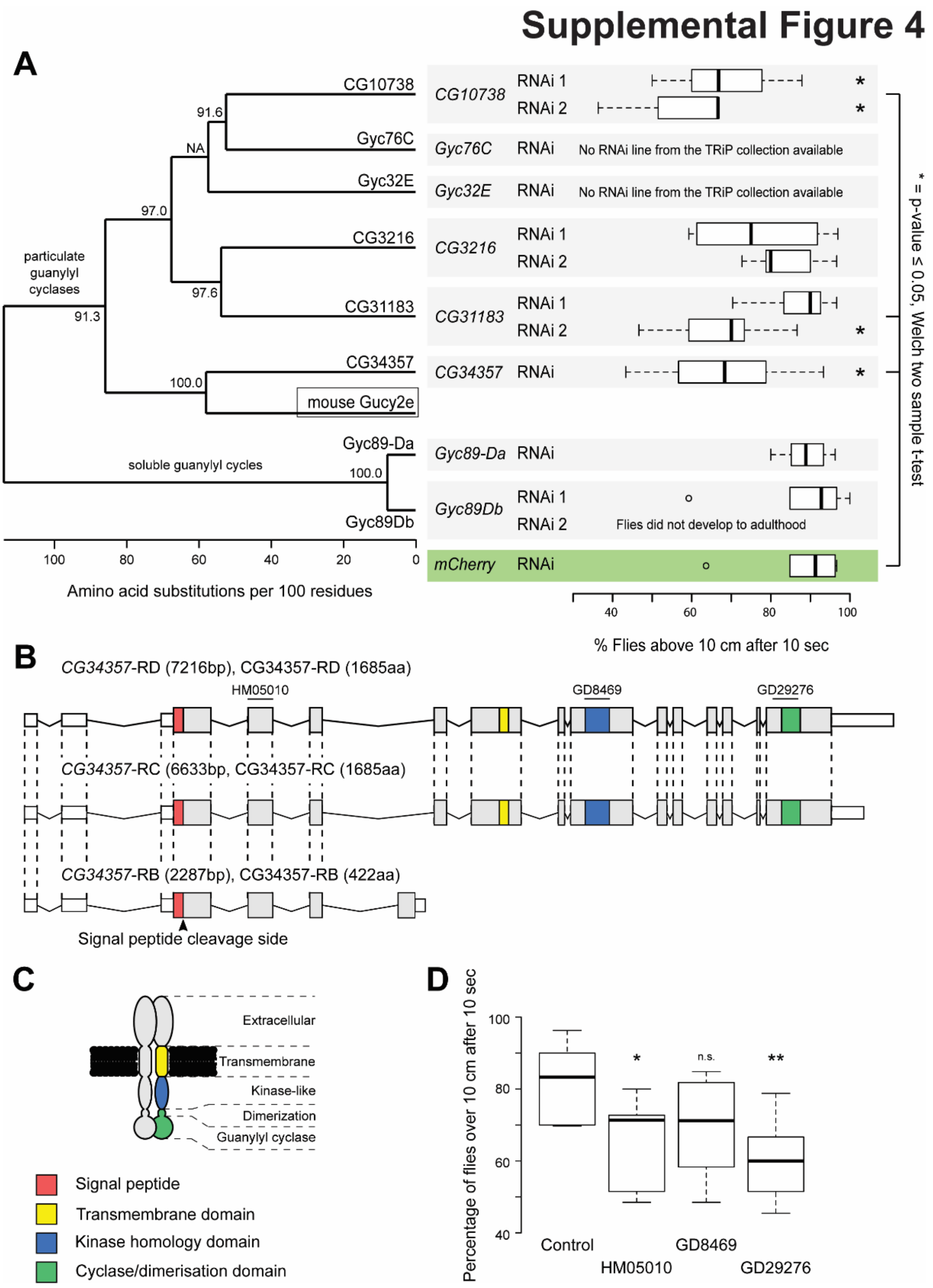


**Supplemental Figure 4: CG34357 is a homologue of mouse Gucy2e and its downregulation in the cholinergic neurons leads to climbing defects.** (**A**) PSI-BLAST search was performed against the fly genome using the mouse Gucy2e protein sequence. All six particulate guanylyl cyclases encoded in the *Drosophila* genome were found in the search (Morton, 2004). The similarity of the protein sequences to each other is shown on the left side (maximum-likelihood phylogenetic tree). Two soluble guanylyl cyclases were included as outgroups. A behavioural screen was conducted to determine if any of the identified enzymes could be involved in climbing (or gravity sensing), and the results are shown on the right side. The genes were knocked down using RNAi; a hairpin against *mCherry* was used as a negative control. The RNAi was driven using the cholinergic-neuronal driver (Gal4$^{Chat19b}$), and flies were raised at 29°C to increase the potential phenotype, as Gal4 is sensitive to temperature. Only fly lines from the TRiP (Transgenic RNAi Project) library were considered (see Methods). When possible, two to three RNAi lines were used against the same gene. Each boxplot corresponds to 60 flies measured in groups of ten flies each. (**B**) *CG34357* gene has three annotated transcripts in the fly. Isoform RC and RD lead to the same protein and only vary in the length of the 3'-UTR. Isoform RB results from one alternative splice event leading to a drastically shorter amino acid sequence containing only extracellular parts of the protein. The different functional domains are colour-coded (legends are described in C). The exons targeted by the hairpins used in D are indicated in the scheme. (**C**) Protein model is drawn analogously to vertebrate particulate guanylyl cyclase (Kuhn, 2016), and functional domains are indicated using the same colours as in B. The signal peptide is not shown in the protein model because it should be cleaved off according to signal peptide prediction (Nielsen, 2017). (**D**) A climbing assay was used to determine the role of *CG34357* in gravity sensing. An RNAi line against *mCherry* was used as a negative control. All RNAis were driven using the Gal4$^{Chat19b}$, and the flies were raised at 29°C to increase the potential phenotype. Each boxplot corresponds to a total of 60 flies measured in sets of ten flies each. In A and D, the significance is determined using the Welch two sample t-test (*p-value < 0.05, **p-value < 0.01).

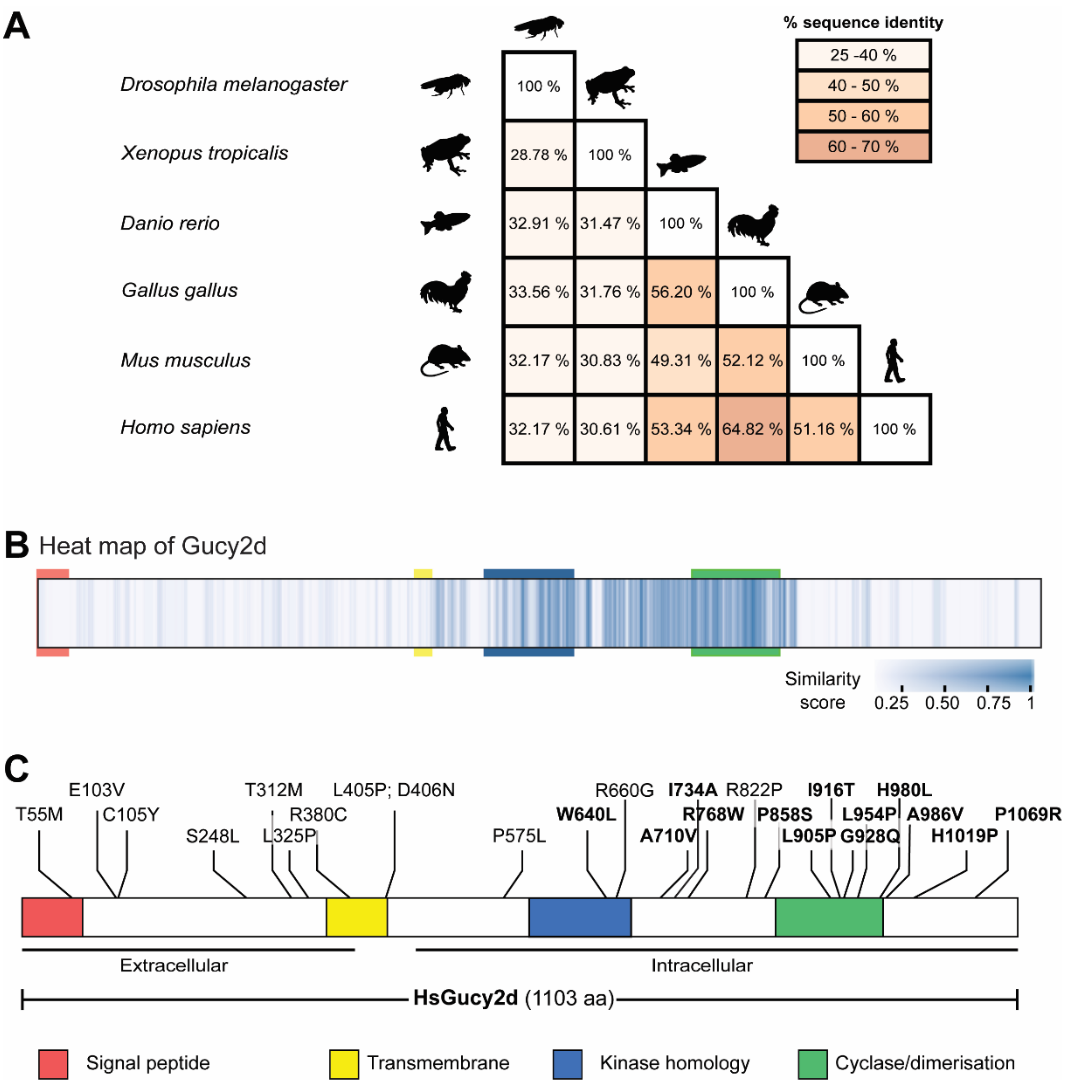


**Supplemental Figure 5: Cross species comparison of Gucy2d homologues and overview of known mutations in human Gucy2d in patients with Leber congenital amaurosis (LCA).** (**A**) Matrix shows pair-wise comparison of different Gucy2d homologues between different vertebrate species and *Drosophila melanogaster*. Numbers indicate percentages of identical residues in pair-wise alignments. Even within vertebrate species the sequence identity is rather low. (**B**) Multiple sequence alignment of Gucy2d from twelve species is represented as a heat map generated using JProfileGrid2 (a list of the species is available in Supplemental Table 3). Each position of the alignment is shown as a box, which is then colour-coded according to the similarity score. The relative positions of the four critical domains of Gucy2d are indicated with the specific-coloured boxes. (**C**) Known sites mutated in patients with LCA. Residues highlighted in bold are conserved in *Drosophila melanogaster* (Supplemental Table 08). There is no clear hotspot region for mutations in humans (de Castro-Miro et al., 2014; Feng et al., 2020; Jacobson et al., 2013; Li et al., 2011; Liu et al., 2020; Salehi Chaleshtori et al., 2020; Tucker et al., 2004; Zagel and Koch, 2014).

### Supplemental Tables:

**Supplemental Table 01:** IFT and BBSome homologues in various species. *Drosophila* homologues were annotated according to https://flybase.org/.

| | ***Chlamydomonas rheinhardii*** | ***Homo sapiens*** | ***Drosophila melanogaster*** |
|---|---|---|---|
| IFT-A (7 members) | IFT43 | IFT43 | CG5780 |
| | IFT121 | WDR35 | Oseg4 |
| | IFT122/FAP | WDR10 | Oseg1 |
| | IFT139 | THM1/TTC21B | *Not conserved* |
| | IFT140 | WDTC2 | RempA/Oseg3 |
| | IFT144 | WDR19 | Oseg6 |
| IFT-B (16 members) | IFT20 | IFT20 | CG30441 |
| | IFT22/FAP9 | RabL5 | *Not conserved* |
| | IFT25/FAP232 | IFT25 | *Not conserved* |
| | IFT27 | RabL4 | *Not conserved* |
| | IFT38/FAP22 | Cluap | *Not conserved* |
| | IFT46 | IFT46 | CG15161 |
| | IFT52/Bld1 | IFT52 | Osm-6 |
| | IFT54/FAP116 | TRAF3IP1 | CG3259 |
| | IFT56/DYF-13 | TTC26 | *Not conserved* |
| | IFT57 | Hippi | Che-13 |
| | IFT70/FAP259 | TTC30A/B | *Not conserved* |
| | IFT74 | IFT74 | *Not conserved* |
| | IFT80 | IFT80 | Oseg5 |
| | IFT81 | IFT81 | *Not conserved* |
| | IFT88 | IFT88 | NompB |
| | IFT172 | IFT172 | Oseg2 |
| BBSome (10 members) | BBS1 | BBS1 | BBS1 |
| | BBS2 | BBS2 | *Not conserved* |
| | BBS3 | BBS3 | Arl6 |
| | BBS4 | BBS4 | BBS4 |
| | BBS5 | BBS5 | BBS5 |
| | BBS6 | BBS6 | *Not conserved* |
| | BBS7 | BBS7 | *Not conserved* |
| | BBS8 | BBS8 | BBS8 |
| | BBS9 | BBS9 | BBS9 |
| | BBS18 | BBSIP10 | *Not conserved* |

**Supplemental Table 02**: IFT88 protein sequences used for comparison.

| **Species** | **Abbreviations** | **Accession number** |
|---|---|---|
| ***Aedes aegypti*** | Ae | EAT40549 |
| ***Anopheles gambiae*** | Ag | XP_556988.3 |
| ***Danio rerio*** | Dr | XP_005167586 |

| | | |
|---|---|---|
| ***Drosophila erecta*** | De | XP_001973983.2 |
| ***Drosophila melanogaster*** | Dm | NP_724347.3 |
| ***Drosophila simulans*** | Ds | EDX05743.1 |
| ***Gallus gallus*** | Gg | XP_015134793.1 |
| ***Homo sapiens*** | Hs | NP_001305422 |
| ***Mus musculus*** | Mm | NP_033402 |
| ***Tribolium castaneum*** | Tc | EFA00677 |
| ***Xenopus tropicalis*** | Xt | F6W448 |

**Supplemental Table 03**: Protein sequences used for bioinformatic analysis on guanylyl cyclases. Whenever several isoforms were available for the same gene, the longest protein sequence was chosen.

| **Name** | **Species** | **Accession number** |
|---|---|---|
| **Hs_Gucy2d** | *Homo sapiens* | NP_000171 |
| **Mm_Gucy2e** | *Mus musculus* | NP_032218 |
| **Gg_retGC1** | *Gallus gallus* | AAC24500 |
| **Dr_retGC2** | *Danio rerio* | NP_001103165 |
| **Xt_retGC1** | *Xenopus tropicalis* | XP_002942678 |
| **Ms_GC-II** | *Manduca sexta* | AAN16469.1 |
| **Tc_retGC2** | *Tribolium castaneum* | KYB27701.1 |
| **Ag_AAEL007359-PA** | *Aedes aegypti* | XP_001658332.1 |
| **Ae_AGAP002233-PA** | *Anopheles gambiae* | XP_307952.5 |
| **Dm_CG10738** | *Drosophila melanogaster* | NP_729905.2 |
| **Dm_CG31183** | *Drosophila melanogaster* | NP_001287342.1 |
| **Dm_CG3216** | *Drosophila melanogaster* | NP_726013 |
| **Dm_CG34357** | *Drosophila melanogaster* | NP_001189166.1 |
| **Dm_Gyc32E** | *Drosophila melanogaster* | NP_001097148.1 |
| **Dm_Gyc76C** | *Drosophila melanogaster* | NP_001163473.1 |
| **Dm_Gyc89-Da** | *Drosophila melanogaster* | NP_001036718.1 |
| **Dm_Gyc89-Db** | *Drosophila melanogaster* | NP_650551.1 |
| **De_GG11511** | *Drosophila erecta* | XP_001978835 |
| **Ds_GD19673** | *Drosophila simulans* | XP_016033187 |

**Supplemental Table 04**: Overview of IFT88 velocities in mouse, *C. elegans* and *Drosophila* (including the results presented in this paper).

| **Species** | **Anterograde velocity** | **Retrograde velocity** | **Reference** |
|---|---|---|---|
| ***Mus musculus* (IMCD3 cells)** | 0.32 µm/s | 0.64 µm/s | (Besschetnova et al., 2010) |

| | | | |
|---|---|---|---|
| ***Mus musculus* (IMCD3 cells)** | 1.19 µm/s | 1.0 µm/s | (Ishikawa et al., 2014) |
| ***Mus musculus* (IMCD3 cells)** | 0.68 µm/s | 0.35 µm/s | (Ye et al., 2013) |
| ***Mus musculus* (IMCD3 cells)** | 0.38 µm/s | 0.56 µm/s | (Tran et al., 2008) |
| ***Mus musculus* (MEF)** | 1.09 µm/s | 1.06 µm/s | (He et al., 2014) |
| ***Mus musculus (olfactory neurons)*** | 0.23 µm/s | 0.14 µm/s | (Williams et al., 2014) |
| ***C. elegans*** | 0.71 µm/s (middle segment); 1.31 µm/s (distal segment) | 1.22 µm/s | (Wei et al., 2012) |
| ***Drosophila melanogaster*** | 0.44 µm/s (proximal and distal segment) | 0.12 µm/s (proximal segment); 0.77 µm/s (outer segment) | (Lee et al., 2018) |
| ***Drosophila melanogaster*** | 0.44 µm/s (proximal segment) | 2.09 µm/s (proximal segment) | This study |

**Supplemental Table 05**: Details of the antibodies used in this paper.

| **Antibody** | **Dilution** | **Species** | **Source** |
|---|---|---|---|
| **Primary antibodies** | | | |
| anti-GFP | 1:1000 | rabbit | Invitrogen, USA |
| anti-GFP | 1:1000 | chicken | Aves, USA |
| anti-GFP | 1:1000 | rabbit | Roche, Germany |
| anti-GFP | 1:1000 | rabbit | Abcam, UK |
| anti-HA | 1:1000 | mouse | Biolegend, UK |
| anti-PLP | 1:1000 | chicken | (Fu and Glover, 2012) |
| anti-Glutamylated tubulin GT335 | 1:500 | mouse | (Wolff et al., 1992) |
| anti-NompC | 1:200 | rabbit | (Cheng et al., 2010) |
| anti-Iav | 1:500 | rat | (Gong et al., 2004) |
| anti-acetylated tubulin | 1:500 | mouse | Sigma, USA |
| **Secondary antibodies** | | | |
| anti-chicken FITC | 1:500 | donkey | Jackson Labs, USA |

| | | | |
|---|---|---|---|
| anti-rat Rhodamine | 1:500 | donkey | Jackson Labs, USA |
| anti-mouse IRDye 680 | 1:10000 | goat | LI-COR, USA |
| anti-mouse IRDye 800 | 1:10000 | goat | LI-COR, USA |
| anti-rabbit IRDye 800 | 1:10000 | goat | LI-COR, USA |
| anti-rabbit Alexa647 | 1:10000 | donkey | Jackson Labs, USA |
| anti-rabbit Alexa488 | 1:500 | goat | Molecular Probes, USA |
| anti-rabbit Alexa647 | 1:500 | donkey | Jackson Labs, USA |
| anti-chicken Rhodamine | 1:500 | donkey | Jackson Labs, USA |
| anti-rat Rhodamine | 1:500 | goat | Jackson Labs, USA |
| anti-mouse Alexa488 | 1:500 | goat | Molecular Probes, USA |
| anti-mouse Alexa647 | 1:500 | goat | Life technologies, USA |

**Supplemental Table 06**: Primers used for the experiments in this paper and the annealing temperatures ($T_a$) used. The OligoCalc tool was used for prediction (Kibbe, 2007); extension times were chosen according to the length of expected product and the properties of the respective polymerase according to the manufacturer.

| **Name** | **Purpose** | **Sequence** | **$T_a$** |
|---|---|---|---|
| **Dm*ift88*_FL_gateway_fwd** | Gateway cloning Dm*ift88* | GGGACAAGTTTGTACA AAAAAGCAGGCTTCAT GACTTCTCAAATAACT GCTAACGGAACGC | 52°C |
| **Dm *ift88*_FLS_gateway_rev** | Gateway cloning Dm*ift88* (with Stop codon) | GGGGACCACTTTGTAC AAGAAAGCTGGGTCTC AAATAGGCAATAAGCT TTCGGG | 52°C |
| **Dm *ift88*_seq_fwd** | For sequencing Dm*ift88* coding sequence | GCAGCCACAGTGAAACATCG | -- |
| **Dm *ift88*_seq_rev** | For sequencing Dm*ift88* coding sequence | AACTCAGGTTGGTCAGAGCG | -- |
| **Dm *ift88*_seq_rev2** | For sequencing Dm*ift88* coding sequence | AGACGGGTGTATTCCGATGC | -- |
| **Dm *ift88*-RD_RT-PCR_fwd** | Quantify mRNA expression of Dm*ift88*-RD isoform | CCCCTACACGTCCATTCTGC | 54°C |
| **Dm *ift88*-RD_RT-PCR_rev** | Quantify mRNA expression of | CCCCTACACGTCCATTCTGC | 54°C |

| | | | |
|---|---|---|---|
| | Dm*ift88*-RD isoform | | |
| **Dm *ift88*_RC_RT-PCR_fwd** | Quantify mRNA expression of Dm*ift88*-RC isoform | CGCTGGAATTAGGGGACCTG | 54°C |
| **Dm *ift88*-RC_RT-PCR_rev2** | Quantify mRNA expression of Dm*ift88*-RC isoform | AGGATCAAACTGCAGAAATCG | 54°C |
| ***elF*-1A_RT-PCR_fwd** | Quantify mRNA expression of *elF1A* (house keeping gene) | GATATACTGGTTCCCCGCGA | 54°C |
| ***elF*-1A_RT-PCR_rev** | Quantify mRNA expression of *elF1A* (house keeping gene) | GGCTTGTTGGCGACCAATTTT | 54°C |
| ***Su(Tpl)*_RT-PCR_fwd** | Quantify mRNA expression of *Su(Tpl)* (house keeping gene) | AGCCACAAATCCATGCAGAG | 54°C |
| ***Su(Tpl)*_RT-PCR_rev** | Quantify mRNA expression of *Su(Tpl)* (house keeping gene) | TGGACGTTGACTTCTTGTTGT | 54°C |
| **TBP_RT-PCR_fwd** | Quantify mRNA expression of TBP (house keeping gene) | TTATGCGAATCCGAGAGCCC | 54°C |
| **TBP_RT-PCR_rev** | Quantify mRNA expression of TBP (house keeping gene) | GTCGAGGAACTTTGCAGGGA | 54°C |
| **Dm*Gucy2d* (*CG34356*)_5UTR_fwd** | Outer primer pair for nested PCR strategy to amplify FL-Dm*Gucy2d* coding sequence | GGCAATGTTGGAAGGCACTG | 52°C |
| **Dm*Gucy2d*_3UTR_rev** | See above | CCCATGTCGTCAGCCATCTAA | |
| **Dm*Gucy2d*_gateway_FLfwd** | Gateway cloning of FL Dm*Gucy2d* | GGGGACAAGTTTGTACAAAAAAGCAGGCTTCATGAAGTTAACAACCTGTCAAATTGCTAAAG | 53°C |
| **Dm*Gucy2d*_gateway_FLrev** | Gateway cloning of FL Dm*Gucy2d* | GGGGACCACTTTGTACAAGAAAGCTGGGTCTCTGCTCAGCAGTTGCTCTGCATTAAGGG | 53°C |
| **Dm*Gucy2d*_trunc_fwd** | Gateway cloning of T1-Dm*Gucy2d* | GGGGACAAGTTTGTACAAAAAAGCAGGCTTCATGCGTAAGCGTCTCTCCAAGGG | 53°C |

| | | | |
|---|---|---|---|
| **Dm*Gucy2d*_trunc_rev1** | Gateway cloning of T2-Dm*Gucy2d* | GGGGACCACTTTGTACAAGAAAGCTGGGTCGTTCGCCGTGGTTGTGATG | 53°C |
| **Dm*Gucy2d*_trunc_rev2** | Gateway cloning of T3-Dm*Gucy2d* | GGGGACCACTTTGTACAAGAAAGCTGGGTCTCTGGTTCTTGGCGGAAGTC | 53°C |
| **Dm*Gucy2d*_trunc_rev3** | Gateway cloning of T4-Dm*Gucy2d* | GGGGACCACTTTGTACAAGAAAGCTGGGTCAAATTCTTCCGGGTCGACGG | 53°C |
| **Dm*Gucy2d*_trunc_rev4** | Gateway cloning of T5-Dm*Gucy2d* | GGGGACCACTTTGTACAAGAAAGCTGGGTCAGATTCTCCAATAGGCGGCG | 53°C |
| **Dm*Gucy2d*_seq_fwd1** | To confirm PCR products through sequencing | TGGCAGGACAATACGTGACC | -- |
| **Dm*Gucy2d*_seq_fwd2** | To confirm PCR products through sequencing | GGACTGGTCGTTTCGGCTAA | -- |
| **Dm*Gucy2d*_seq_fwd3** | To confirm PCR products through sequencing | TCCTGGCGCATCCATATGTC | -- |
| **Dm*Gucy2d*_seq_rev1** | To confirm PCR products through sequencing | GAACCCATGCTGAGTGTCCA | -- |
| **Dm*Gucy2d*_seq_rev2** | To confirm PCR products through sequencing | TTTCTCGAGGTCAAGGCACC | -- |
| **Dm*Gucy2d*_seq_rev3** | To confirm PCR products through sequencing | GGTGGAATTGGAGTCCAGGG | -- |

**Supplemental Table 07**: Fly stocks used in this study. (BSC – Bloomington Stock Center, VDRC – Vienna Drosophila Resource Center, DGRC – Drosophila Genomics Resource Center)

| **Name** | **Genotype** | **Source** |
|---|---|---|
| **Gal4$^{Tub}$** | y[1] w[*]; P{w[+mC]=tubP-GAL4}LL7/TM3, Sb[1] Ser[1] | BSC, # 5138 |
| ***Tub*-Gal80$^{ts}$** | w[*]; sna[Sco]/CyO; P{w[+mC]=tubP-GAL80[ts]}7 | Made from BSC, #7017 |
| **Gal4$^{Chat19b}$** | w[*]; Chat-Gal4/CyO | (Jana et al., 2011) |
| **Gal4$^{Chat19b}$, UAS-*GFP*** | w[*]; Chat-Gal4, UAS-GFP | (Jana et al., 2011) |
| **Gal4$^{Iav}$** | w[*];; Iav-Gal4 | BSC, # 52273 (a gift from Martin Göpfert's lab) |

| | | |
|---|---|---|
| **40xUAS-IVSmCD8::GFP** | w[*];; P{y[+t7.7] w[+mC]=40XUAS-IVSmCD8::GFP}attP2 | BSC, #32195 |
| **UAS-mCD8::GFP** | w[*]; UAS-mCD8::GFP/CyO.Z; MKRS/TM6B | a gift from Elio Sucena's lab |
| **UAS-GFP::DmIFT88** | W[1];;UAS-GFP::DmIFT88/TM6B | This study (made using IGC Fly transgene facility) |
| **endo-GFP::DmIFT88** | w[*]; endo-GFP::DmIFT88 | (Han et al., 2003) (a gift from Daniel Eberl) |
| ***mCherry* RNAi** | y[1] sc[*] v[1]; P{y[+t7.7] v[+t1.8]=VALIUM20-mCherry}attP2 | BSC, # 35785 |
| **Dm*ift88* RNAi** | y[1] v[1]; P{y[+t7.7] v[+t1.8]=TRiP.JF03080}attP2 | BSC, # 28665 |
| ***CG10738* RNAi 1** | y[1] v[1];; P{y[+t7.7] v[+t1.8]=TRiP.HMS01814}attP2 | BSC, #38346 |
| ***CG10738* RNAi 2** | y[1] v[1];; P{y[+t7.7] v[+t1.8]=TRiP.HM05067}attP2 | BSC, #28580 |
| ***CG31383* RNAi 1** | y[1] v[1];; P{y[+t7.7] v[+t1.8]=TRiP.HM05092}attP2/TM3, Sb[1] | BSC, #28604 |
| ***CG31383* RNAi 2** | y[1] v[1]; P{y[+t7.7] v[+t1.8]=TRiP.HMJ22232}attP40 | BSC, #58224 |
| ***CG3216* RNAi 1** | y[1] sc[*] v[1];; P{y[+t7.7] v[+t1.8]=TRiP.HM05270}attP2/TM3, Sb[1] | BSC, #31877 |
| ***CG3216* RNAi 2** | y[1] v[1];; P{y[+t7.7] v[+t1.8]=TRiP.HMC04174}attP2 | BSC, #55895 |
| **Dm*Gucy2d* (*CG34357*) GD29276** | w1118;; UAS-IR[GD8469] | VDRC, #29276/GD |
| **Dm*Gucy2d* GD8469** | w1118; UAS-IR[GD8469] | VDRC, #8469/GD |
| **Dm*Gucy2d* HM05010** | y[1] v[1];; P{y[+t7.7] v[+t1.8]=TRiP.HM05010}attP2 | BSC, #28524 |
| **Dm*Gucy2d* [NP0270]** | y[*] w[*];; P{w[+mW.hs]=GawB}CG34357[NP0270] /TM6, P{w[-]=UAS-lacZ.UW23-1}UW23-1 | DGRC, #103574 |
| ***Gyc89*-Da RNAi** | y[1] sc[*] v[1];; P{y[+t7.7] v[+t1.8]=TRiP.HM05246}attP2/TM3, Sb[1] | BSC, #30502 |
| ***Gyc89*-Db RNAi 1** | y[1] v[1];; P{y[+t7.7] v[+t1.8]=TRiP.HM05207}attP2 | BSC, #29529 |
| ***Gyc89*-Db RNAi 2** | y[1] v[1]; P{y[+t7.7] v[+t1.8]=TRiP.HMJ22088}attP40 | BSC, #58139 |
| **UAS-DmGucy2d-T1::GFP** | w[*];; UAS- DmGucy2d -T1::GFP/TM6B | This study (made using IGC Fly transgene facility) |

| | | |
|---|---|---|
| **UAS-DmGucy2d::GFP** | w[*]; UAS-DmGucy2d::GFP/CyO | This study (made using BestGene facility) |
| **UAS-DmGucy2d::GFP** | w[*];; UAS-DmGucy2d::GFP/TM3 | This study (made using BestGene facility) |

**Supplemental Table 08**: **Screenshots of alignments of selected sequences of intracellular domains of Gucy2d from diverse species visualised with "Multiple Alignment Show".** Sequences were aligned using MUSCLE (http://www.ebi.ac.uk/Tools/msa/muscle/). In addition to the six species shown in Supplemental Figure 5A, six insects species were included in the analysis to avoid a vertebrate bias in the multiple sequence alignment (Supplemental Table 03). Note that the residues that are mutated in human LCA patients and are evolutionarily conserved are marked with red boxes (de Castro-Miro et al., 2014; Feng et al., 2020; Jacobson et al., 2013; Li et al., 2011; Liu et al., 2020; Salehi Chaleshtori et al., 2020; Tucker et al., 2004; Zagel and Koch, 2014).

**W640L**

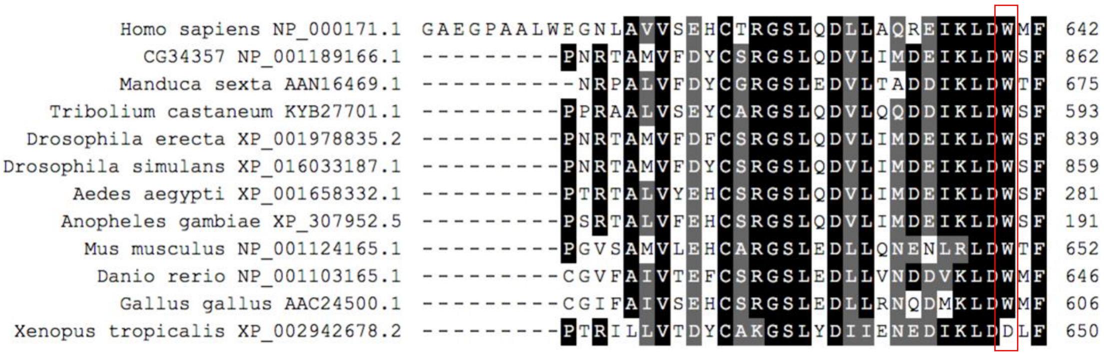


**A710V**

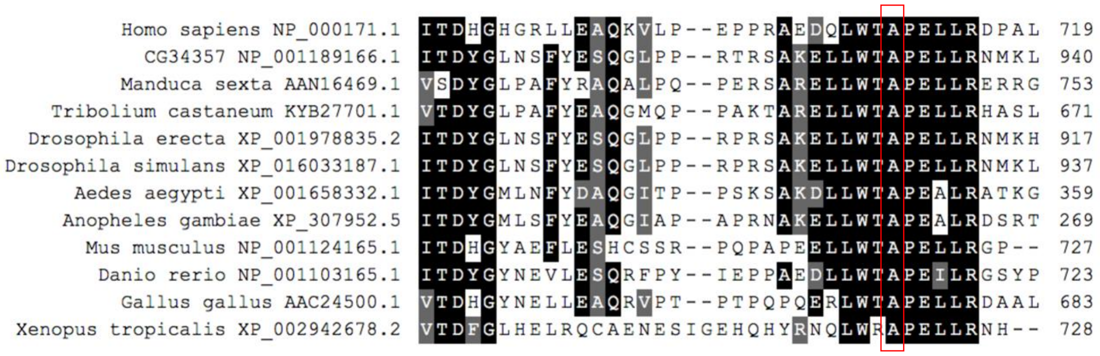

### I734A

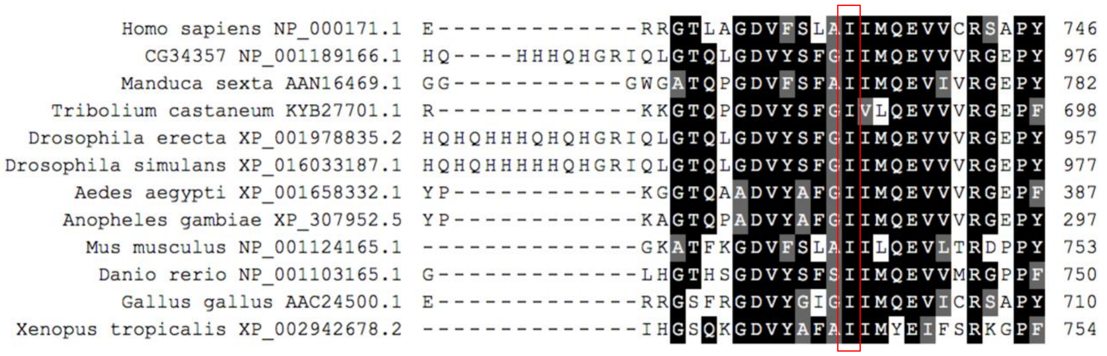


### R768W

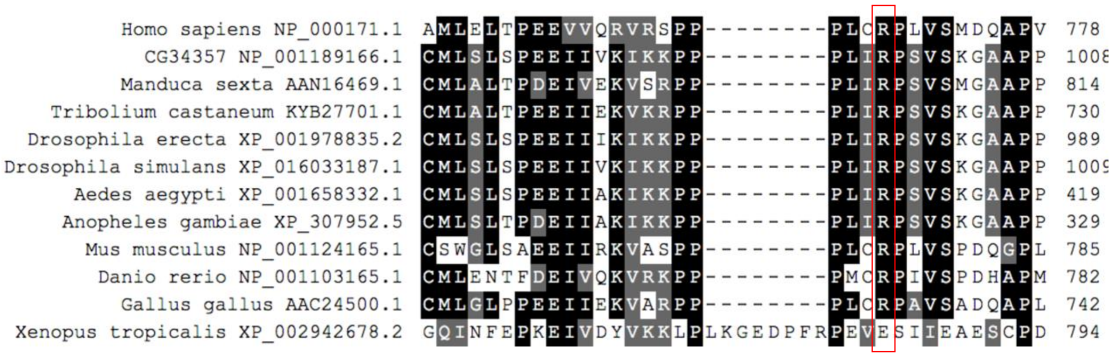


### P858S

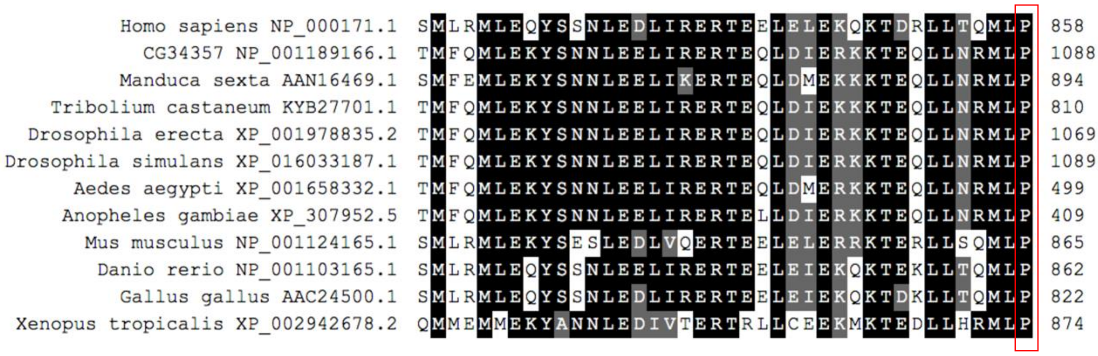


### L905P, I916T, G928Q

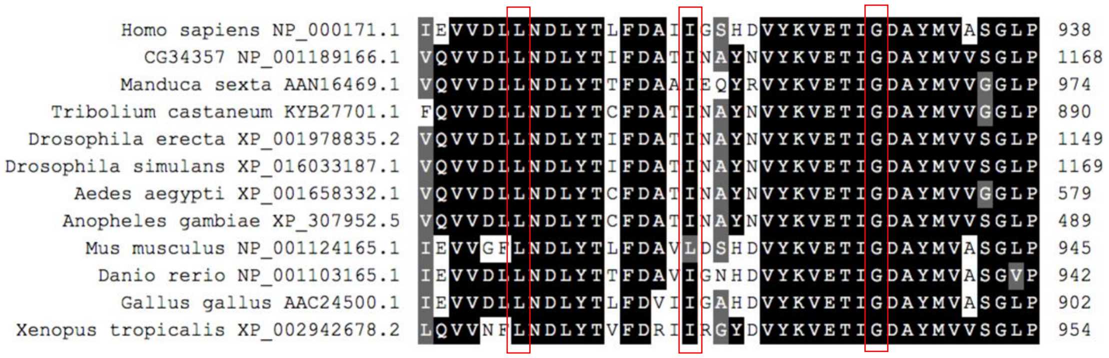

**L954P**

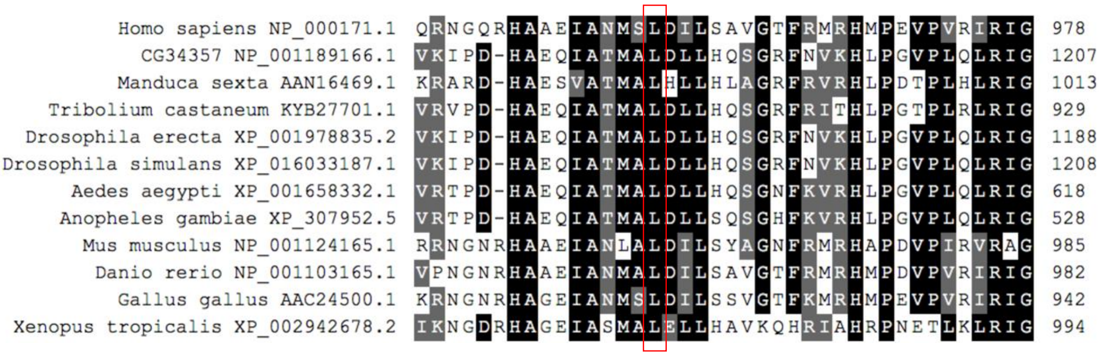


**H980L, A986Vfs*76**

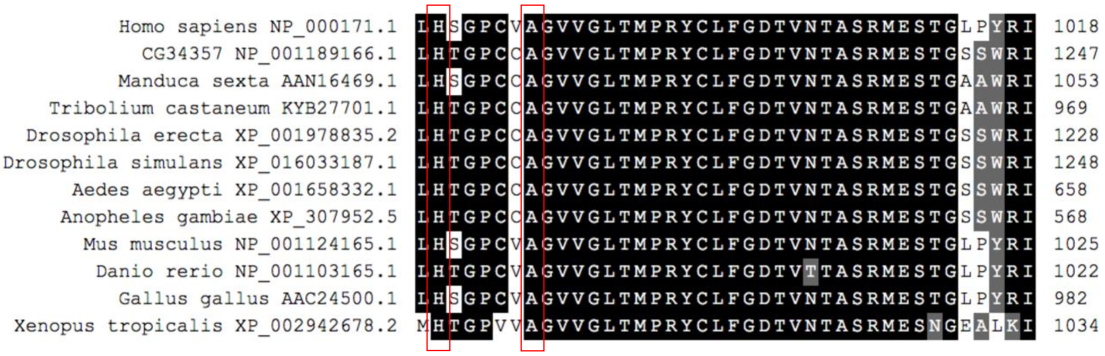


**H1019P**

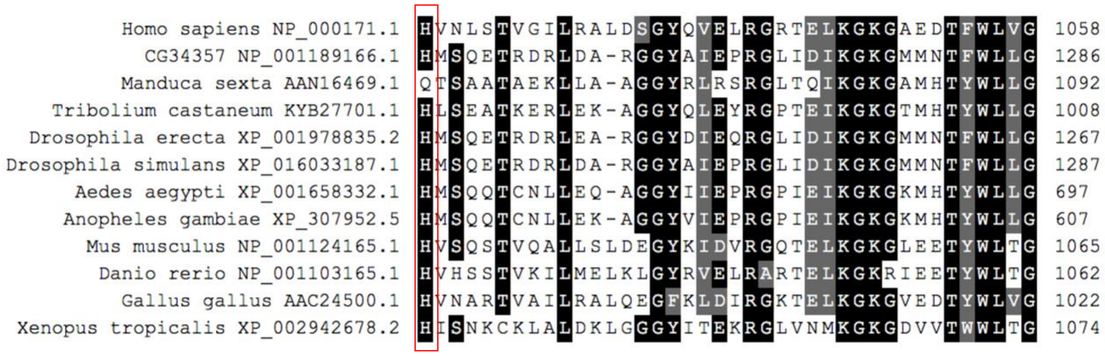


**P1069R**

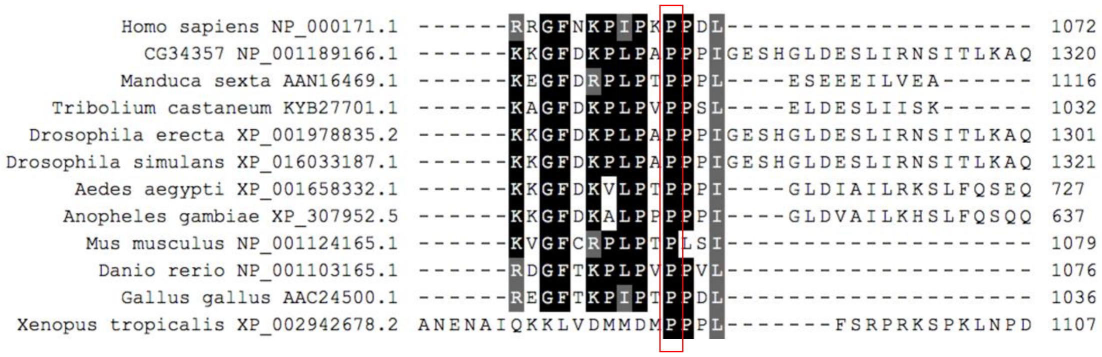

### Materials and methods:

#### Bioinformatics analysis

**For IFT88:** The gene model for Dm*ift88* was extracted from the Ensembl Genome Browser (Yates et al., 2016). The number of tetratricopeptide repeat domains (TPR) was predicted using the TPRpred tool (Karpenahalli et al., 2007). For phylogenetic analysis, IFT88 protein sequences of eleven metazoan species were used (see Supplemental Table 02). Whenever several *ift88* isoforms were reported for one species, the largest one was chosen for analysis. The MegAlign program (Lasergene suite, Version 8.1.3, DNASTAR©) was used to generate a phylogenetic tree summarising sequence similarities as a function of the number of amino acid substitutions between sequences. Sequences were aligned using the ClustalW algorithm with default settings. Bootstrapping analysis was performed with default values to calculate the "support value" of each branching point in the tree (Figure 1Aii left). Multiple sequence alignments are presented as a heatmap. An alignment file in the FASTA format using the web browser-based MUSCLE tool was created (Edgar, 2004) to extract sequence identities. Subsequently, the ProfilGrid tool (Roca et al., 2008) was used with default settings to assign a similarity score to each position in the alignment, with the score assuming values between 0 and 1. The results are presented as a heatmap, which was visualised with RStudio (RStudio®, USA) and the ggplot2 package (Wickham, H. (2016). ggplot2. Use R! doi:10.1007/978-3-319-24277-4) (Figure 1Aii right).

**For guanylyl cyclases, including Dm*Gucy2d*/*CG34357*:** Sequence analysis for guanylyl cyclases (Supplemental Figure 3A) was performed as described above. For the accession number of guanylyl cyclases protein sequences used in this analysis, see Supplemental Table 03. The gene model for *CG34357* was taken from the Ensembl Genome Browser (Yates et al., 2016).

#### Cryosectioning, immunolabeling and image analysis

Histological sections of the adult antenna were prepared as described before (Jana et al, 2016, Mishra, 2015). If not stated otherwise, preparation steps were carried out at room temperature. Flies were anaesthetised using $CO_2$ and decapitated. The heads were collected in pre-cooled fixation buffer (4% paraformaldehyde, 75 mM PIPES buffer (pH 7.6), 0.05% Triton-X, picric acid) and incubated for 40 minutes. Good fixation requires the heads to sink to the bottom of the reaction tube. The heads were washed three times in PBST (PBS with 0.05% Triton-X) and then incubated on a rotator in PBST with 10% sucrose, first for one hour at room temperature and then, overnight, in 25% at 4°C. The heads were embedded and oriented appropriately in OCT (optimal cutting temperature formulation; Tissue-Tek, Sakura Finetek, USA) and the OCT was frozen on dry ice. Head samples were sectioned at 12 µm thickness in a Leica Cryostat CM 3050 S (Leica, Germany). The slices were collected on Poly-lysine coated slides (Sigma, USA).

Sections were washed with PBST (PBS with 0.05% Triton-X) three times for 10 minutes each, followed by incubation in blocking buffer (10% BSA in PBST) for one hour and subsequent incubation with the primary antibody diluted in blocking buffer overnight at 4°C. Samples were washed three times in blocking buffer and incubated for one hour with secondary antibody in blocking buffer. Finally, after three additional washing steps using PBS for 5 minutes each, samples were embedded in Vectashield supplemented with DAPI (Vector Laboratories). 12- bit images were acquired on a TCS SP5 upright confocal microscope (Leica, Germany) using an PL APO CS 63.0x1.40 OIL objective with a pinhole at 1 Airy unit and a pixel size of 80-200 nm (lateral) and 400nm (axial). For details of the antibodies used, see Supplemental Table 05.

To quantify in the DmIFT88 knockdown experiments the signal intensities of Inactive (Iav; a marker for the proximal cilium) antibody staining, and the GFP signal of the T1-DmGucy2d::GFP at the proximal cilium (Figures 2D, 9 days old flies; and Figure 5E, 6 days old flies), image stacks were analyzed with the Imaris software v.9.2.1 (Bitplane). For Figure 2D: The Iav channel was used to generate segmented volume surfaces and the Iav intensity values were retrieved from Imaris, averaged per antennae in Excel (Microsoft Office, USA) and normalised to the negative control of each repeated experiment. For Figure 5D: The Iav channel was used to generate segmented volume surfaces. Next, mean intensity values of Iav antibody staining, and T1-DmGucy2d::GFP at the segmented Iav surfaces, were retrieved from Imaris, averaged per antennae in Excel (Microsoft Office, USA), and statistically analysed (non-parametric Mann-Witney t-test or Welch Two Sample t-test) in Prism (GraphPad Software, USA). Data were represented on box plots (median ± full range, each dot representing individual averaged values) using Prism 5.2.

To quantify the T1-DmGucy2d::GFP intensity at the end of the dendrites in the adult chordotonal neurons (Figure 5F), the image stacks were deconvolved using the Huygens Deconvolution v17.4 software (Scientific Volume Imaging) with an CMLE algorithm followed by image analysis in Imaris (as mentioned above). This deconvolution step facilitated the segmentation of the crowed dendrite population in the adult antennae by increasing the signal-to-noise ratio (SNR). N>10 antenna in all conditions.

**mRNA isolation from adult antenna and reverse transcription analysis**

Between 100 and 200 flies were decapitated per genotype. Heads were collected in Protein LoBind tubes (Eppendorf, USA) on dry ice. 10 to 20 heads were collected at a time to avoid compromising of the tissue due to long exposure to room temperature. Tubes were snap-frozen in liquid nitrogen. Antennae were dissociated from heads through mechanical force using a vortex mixer (three times for ten seconds, cooled in between again on dry ice). To separate antennae from heads, tubes were opened and inverted. Heads will fall out, whereas the majority of the antennae will adhere to the wall of the tubes. Total mRNA was isolated from antennae using the PureLink$^{TM}$ RNA Mini Kit (Ambion life technologies, USA). Antennae were lysed in 250 µl of lysis buffer. The protocol was carried out as described in the accompanying manual of the kit. No DNAse treatment was performed. The mRNA was always converted to cDNA immediately. For cloning, 1 µg of total mRNA was reverse transcribed. The reverse transcription reaction was performed with the Transcriptor First Strand cDNA Synthesis kit (Roche, Germany). For quantifying mRNA levels, 50 to 70 heads were used. For cloning Dm*ift88*, cDNA was generated using poly-dT primers included in the kit. The cDNA samples were stored at −20°C until further use.

**Cloning and generation of transgenic flies**

**For DmIFT88:** The cDNA for cloning Dm*ift88* was obtained from mRNA isolated from antennae of $w^{1118}$ flies. Dm*ift88* (isoform-RD) was amplified using the KOD polymerase kit (EMD Millipore, USA) in 50 µl reaction volume (for PCR primers see Supplemental Table 06). PCR was set up as outlined for the kit. The reaction mix was supplemented with DMSO. The PCR product was purified using the DNA Clean & Concentrator$^{TM}$ kit (Zymo Research, USA) and adenylated at the 5’ ends through incubation with DreamTaq (Thermofisher, USA) using the appropriate buffer and dNTPs at 70°C for one hour. Subsequently, the PCR product was ligated into pGEM-Teasy (Promega, USA) following the manufacturer's instructions and plasmids were confirmed through sequencing (for sequencing primers see Supplemental Table 06). Using the Gateway system, the coding sequence was first cloned into pDONR 221 in a BP-reaction (Invitrogen), following the instructions for the kit, and subsequently transferred to pTHW and pTGW in an LR-reaction (Invitrogen, USA) to fuse the resulting protein to three HA-tags or one GFP-tag on the N-terminus, respectively (for details on pTHW and pTGW see Drosophila Genomic Resource Center, USA). DmIFT88 was tagged at the N-terminus, because the GFP::DmIFT88 fusion protein was previously shown to rescue the ift88/*nompB* mutant phenotype and can hence be considered fully functional (Han et al., 2003).

**For Dm*Gucy2d*/*CG34357* full length (FL) and truncated T1-DmGucy2d GFP constructs:** The *CG34357* coding sequence was cloned from mRNA isolated from antennae of $w^{1118}$ flies as described above. Dm*Gucy2d* could only be amplified from cDNA produced with random hexamer primers (included in the Transcriptor First Strand cDNA Synthesis kit, Roche, Germany). For primer sequences, see Supplemental Table 06. To enhance the chance to amplify the desired target (coding sequence of *CG34357*-RD, Supplemental Figure 4B) even further, the mRNA was reverse-transcribed using a primer on the 3'-untranslated region of Dm*Gucy2d* (*CG34357*_3UTR_rev 2 µM in a 20 µl reaction volume, 1 µg of total mRNA were used as template). Two rounds of PCRs were performed to obtain the Dm*Gucy2d* with the appropriate overhangs for Gateway cloning (Invitrogen, USA). In the first round the Dm*Gucy2d* coding sequence was amplified using primers located on each of the untranslated regions. For the second PCR, 2 µl of unpurified PCR product of the first PCR was used as a template. Primers containing appropriate sequences for gateway cloning were used for the second PCR. The coding sequence was cloned without the stop codon to allow for gene fusion at the C-terminus. Tagging at the N-terminus was not considered, since the protein is predicted to undergo cleavage of the signal peptide (30 amino acids, Supplemental Figure 4B, arrowhead). The coding sequence was confirmed through sequencing and sub-cloned into pTWG (Drosophila Genomic Resource Centre) for C-terminal fusion with GFP.

Plasmid DNA was amplified in *E. coli* DH5α and purified using ZR Plasmid MiniprepTM-Classic (Zymo research, USA). ZymoPURE$^{TM}$ Midiprep Kit (Zymo research, USA) was used for large scale DNA purification that is required (Dm*ift88*, Dm*Gucy2d* or T1-Dm*Gucy2d* in pTWG) for transfection experiments and fly transgenesis. Injections into *Drosophila* embryos and selection of positive transformants were outsourced (BestGene, USA or IGC transgenics facility, Portugal).

**Detection of Dm*Gucy2d* gene expression:** In order to detect expression of Dm*Gucy2d* in different developmental stages, an enhancer Gal4-trap line (*CG34357*[NP0270]) was obtained (Brand and Perrimon, 1993). Flies carrying the enhancer trap insertion were crossed with flies encoding 40xUAS-IVS-mCD8::GFP. The mCD8::GFP fusion protein is targeted to the membrane of the cells and was used to visualize the morphology of cells. 40 UAS tandem insertions were chosen to increase the signal intensity.

For imaging adult antenna, the cuticle of the heads was cleared as further explained below, and the second antenna segment was imaged entirely on an SP5 Live Upright microscope (Leica, Germany). Confocal Z-stack images were deconvolved using the Huygens v17.4 software (SVI, Netherlands) and 3D reconstructed using the Imaris v6.4 software (Bitplane, USA), which was also used to prepare the movie (Supplemental movie 02). Wandering L3 larvae were mounted and imaged as described for the live-imaging of L3 larvae below.

**Live imaging of chordotonal cilia in L3 larvae**

GFP::Dm*ift88* was expressed in chordotonal neurons using Gal4$^{lav}$ (Gong et al., 2004b). Flies were reared at 25°C to the third larval stage and all experiments were performed with wandering L3 larvae. Larvae were collected and briefly washed in PBS. For imaging, larvae were immobilised between a coverslip and a slide in a drop of PBS (Zhang et al., 2013). The cover slip was held in place using transparent double-sticky tape. Imaging was performed using a Roper Spinning Disc microscope (Nikon). Samples were kept at 25°C through the imaging process. To prevent artefacts from cell death, larvae were always imaged immediately after immobilisation and for a maximum duration of two minutes. Images were acquired every 120 ms with 100 ms exposure time. Images were analysed using the FIJI software (Schindelin et al., 2012) and kymographs were generated using the KymographClear plugin (Mangeol et al., 2016). Data plots were generated using RStudio (RStudio®, USA).

**Quantification of DmIFT88 expression levels from adult antennae**

For measuring Dm*ift88* expression levels, the total mRNA was isolated from antennae of adult flies with specific genotypes up to four days after eclosure using Real-time PCR (RT-PCR). The web browser-based Primer-BLAST tool (Ye et al., 2012) was used to design primers specific for a given transcript. At least one primer in a pair was designed to span an exon-exon junction in order to avoid amplification of genomic DNA that might contaminate the samples. All primers (see Supplemental Table 06) used in this study were tested to only yield a single product and to not amplify genomic DNA. The Dm*ift88* primers were confirmed to only amplify the respective isoform using cDNA (data not shown). 150 ng of total mRNA were reverse transcribed to cDNA in a reaction volume of 20 µl. The cDNA mix was diluted 1:10, and 4 µl of the dilutions were used in each reaction. iTaq™ Universal SYBR® Green Supermix (Bioradm USA) was used to detect amplification of the PCR products. The reactions were set-up up according to the manufacturers protocol and run on the CFX384 Touch™ Real-Time PCR Detection System (Biorad, USA). The total mRNA was isolated three times per genotype and each sample was measured in triplicates. Results were analysed using RStudio (Rstudio®, USA).

**Sound evoked neuronal responses**

Recordings of sound-evoked compound action potentials of chordotonal neurons in the fly's 2nd antennal segment were performed as previously reported (Bechstedt et al., 2010). Measurements were carried out using flies up to six days after eclosure. Action potentials were recorded from head-fixed flies. For recording, a tungsten needle was inserted between head and antenna. The other antenna was immobilised, and the reference electrode was positioned in the thorax. To determine maximum compound action potential amplitudes, flies were stimulated with pure tones at the individual best frequency of their antenna, which was determined from the velocity spectrum of the antenna's mechanical free fluctuations measured with a scanning laser Doppler vibrometer (PSV-400, Polytec, Waldbronn, Germany) (for details, see (Gopfert et al., 2005)). Data was analysed and graphs were generated using RStudio (RStudio®, USA).

**Sample preparation and image acquisition in Transmission Electron Microscope**

The samples for electron microscopy were processed and sectioned as previously described (Jana et al., 2016). Antennae were dissected from the heads and incubated in fixative (2% formaldehyde, 2.5% glutaraldehyde in 0.1 M sodium biphosphate buffer pH 7.2–7.4) overnight. Samples were washed several times in PBS and then incubated for 1.5 hours in 1% osmium tetroxide for postfixation. Samples were washed several times in deionised water before incubating them in 2% uranyl acetate for 20 minutes on a rotator for *en bloc* staining. After this step, samples were dehydrated in a graded sequence of alcohol dilutions (from 50% to a 100%). Ethanol was removed through two incubation steps with propylene oxide for 30 min. Samples were embedded in resin overnight and transferred to a fresh batch of resin for 1h. Under a dissection microscope, the second segment of the antenna was separated from the third. The second segment was placed in the desired orientation into a mold with resin and polymerised. Samples were sectioned ~70 nm thick, collected on formvar-coated copper slot grids and stained with 2% uranyl acetate followed by lead citrate staining. Finally, samples were air-dried and investigated and photographed at 100kV using the Hitachi H-7650 transmission electron microscope.

**Conditional knockdown in the flies**

The Gal4$^{Chat19b}$ driver is active in cholinergic neurons, including chordotonal and olfactory neurons, in all developmental stages of the fly (Jana et al., 2011; Salvaterra and Kitamoto, 2001). Thus, to avoid confounding effects from the ciliogenesis defects and be able to spatiotemporally knockdown our gene of interest in the adults, we adopted a Gal4$^{Chat19b}$-*Tub*Gal80$^{ts}$-UAS-RNAi-based system. The Gal4 activity was repressed through co-expression of a temperature sensitive version of Gal80 under *Tubulin* promoter (denoted as *Tub*Gal80$^{ts}$) (McGuire et al., 2003). At the permissive temperature (18°C) TubGal80ts is functional, acting as a negative regulator by binding to Gal4 and preventing it from attracting polymerase to a UAS element in the fly genome (del Valle Rodriguez et al., 2011). Thus, a candidate (i.e. Dm*ift88*, Dm*Gucy2d*) mRNA will only be reduced through RNAi if flies are kept at the restrictive temperature (29°C). Flies were reared at 18°C, shifted to the restrictive temperature after eclosure (when ciliogenesis is finished), and then submitted to the climbing (negative-gravitaxis) assay for up to two weeks (also see, Figure 2A).

**Climbing (or negative-gravitaxis) assay**

To quantify climbing behaviour, a behavioural readout of sensory function, flies were collected in 24-hour time windows and kept in groups of the same age. The animals were transferred to vials with fresh food every three days. Prior to the behaviour experiments, flies were separated according to sex into groups of ten animals that would be tested jointly. Flies were briefly anesthetised using $CO_2$ for sorting. The flies were kept in the room where the behavioural experiments were conducted for about one hour prior to the actual experiment. This time period was given to allow the animals to acclimatise to potentially different light and temperature regimes. Experiments were conducted in measuring cylinders made of glass. Animals were placed in cylinders a couple of minutes before the assay was started. Cylinders were sealed on top with perforated parafilm to prevent the flies from escaping. Cylinders were tapped until all flies were at the bottom of the cylinder and flies were subsequently allowed to walk up. All experiments were videotaped for one minute and analysed blindly only once a full data set (including control flies) was obtained. The metric used for analysis was the number of flies above ten centimetres after ten seconds, as, in the set-up. The flies expressing a hairpin against *mCherry* (UAS-*mCherry*-IR; negative control) in cholinergic neurons need on average ten seconds to perform this task. Videos were analysed for the whole duration of ten seconds after tapping the cylinder, to rule out that flies reached the ten-centimetre mark and walked back down again. Data was analysed and graphs were generated using RStudio (RStudio®, USA).

**Cell culture and immunoprecipitation experiments**

For co-immunoprecipitation of ectopically expressed proteins, *Drosophila melanogaster* (Dmel) cultured cells were co-transfected with the corresponding constructs (GFP tagged at the C-terminus of either DmGucy2d full-length or fragments (T1-T5)) with 3xHA::DmIFT88. The co-overexpression of GFP alone with 3xHA::DmIFT88 serves as a negative control in these experiments. Cells were seeded at a density of $3x10^6$ cells/well in 6-well plates (Orange Scientific). One hour later they were transfected using Effectene Transfection Reagent (Quiagen) according to manufacturer (see Supplementary Table 06 for primers information). Cells were harvested three days after transfection. Cells were resuspended, the suspension was collected and centrifuged and pellets were stored at -80°C, if not processed immediately. Ectopically expressed DmGucy2d::GFP was immunoprecipitated at 4ºC for 2 hrs from Dmel cell lysates (three wells in total for each condition per experiment) using polyclonal anti-GFP antibodies and protein A or G magnetic Dynabeads (Thermo Fisher Scientific). 2 µg anti-GFP antibodies were added to protein-A or -G magnetic beads and incubated for 30 min at room temperature. Cells were homogenised in lysis buffer: 50 mM Tris-HCl pH 8, 250 mM NaCl, 1 mM DTT, 2% NP-40, 0.5% of SDS, 0.5% sodiumdeoxycholate, 1× protease inhibitor, 5yg/ml Leupeptin and 15yg/ml Aprotinin, 0.1% Digitonin at 4 C for 30 min. Then, benzonase was added to the lysate (final concentration 0.25 units/µl) and incubated for another 15 min. After centrifugation at 13,500 g for 20 min at 4ºC, the pre-cleared supernatants were incubated with the coupled beads with antibodies. After several washes with lysis buffer, bead pellets were boiled in SDS sample buffer, separated by 4-15 % gradient SDS-PAGE and transferred onto LI-COR nitrocellulose membranes for Odyssey. GFP, DmGucy2d::GFP constructs and HA::DmIFT88 were visualised with anti-GFP or anti-HA antibodies. Secondary LI-COR antibodies IRDye 680RD over IRDye 800 (Li-COR) were used as a second step (see Supplementary Table 05 for details on the antibodies used).

**Cuticle clearing and imaging of antennae whole-mounts**

The cuticle of the adult flies is opaque and autofluorescent, which makes this organ generally unsuitable for imaging. A clearing method that made the cuticle transparent without affecting the morphology of the cells/tissues was developed to image the cytoplasmic GFP expressed in the chordotonal neurons in the adult flies with different genotypes. It allowed imaging of the whole antenna without mechanical manipulation. The whole heads were fixed in 4 % of paraformaldehyde in PBS (supplemented with 0.05 % Triton-X) for 40 minutes on ice (Kolesova et al., 2016). Heads were then transferred to a reaction tube containing FocusClear (CelExplorer, Taiwan) and incubated overnight. For imaging, heads were mounted on a slide in MountClear (CelExplorer, Taiwan). Several layers of clear tape were used as spacer between slide and coverslip. Images were acquired on a SP5 Upright microscope (Leica, Germany). The size of the image stack was chosen to contain the whole antenna segment. Images were then the analysed using ImageJ or Imaris v6.4 software (Bitplane).

**Statistical analysis**

All statistical analyses were performed using either RStudio (RStudio®, USA) or Prism (GraphPad Software, USA). The details of the statistical methods used to analyse a given set of experiments are mentioned in the relevant figure legends.

### References:

Bechstedt, S., J.T. Albert, D.P. Kreil, T. Muller-Reichert, M.C. Gopfert, and J. Howard. 2010. A doublecortin containing microtubule-associated protein is implicated in mechanotransduction in Drosophila sensory cilia. *Nature communications*. 1:11.

Besschetnova, T.Y., E. Kolpakova-Hart, Y. Guan, J. Zhou, B.R. Olsen, and J.V. Shah. 2010. Identification of signaling pathways regulating primary cilium length and flow-mediated adaptation. *Current biology : CB*. 20:182-187.

Cheng, L.E., W. Song, L.L. Looger, L.Y. Jan, and Y.N. Jan. 2010. The role of the TRP channel NompC in Drosophila larval and adult locomotion. *Neuron*. 67:373-380.

de Castro-Miro, M., E. Pomares, L. Lores-Motta, R. Tonda, J. Dopazo, G. Marfany, and R. Gonzalez-Duarte. 2014. Combined genetic and high-throughput strategies for molecular diagnosis of inherited retinal dystrophies. *PloS one*. 9:e88410.

del Valle Rodriguez, A., D. Didiano, and C. Desplan. 2011. Power tools for gene expression and clonal analysis in Drosophila. *Nature methods*. 9:47-55.

Edgar, R.C. 2004. MUSCLE: multiple sequence alignment with high accuracy and high throughput. *Nucleic acids research*. 32:1792-1797.

Feng, X., T. Wei, J. Sun, Y. Luo, Y. Huo, P. Yu, J. Chen, X. Wei, M. Qi, and Y. Ye. 2020. The pathogenicity of novel GUCY2D mutations in Leber congenital amaurosis 1 assessed by HPLC-MS/MS. *PloS one*. 15:e0231115.

Fu, J., and D.M. Glover. 2012. Structured illumination of the interface between centriole and peri-centriolar material. *Open biology*. 2:120104.

Gong, Z., W. Son, Y.D. Chung, J. Kim, D.W. Shin, C.A. McClung, Y. Lee, H.W. Lee, D.J. Chang, B.K. Kaang, H. Cho, U. Oh, J. Hirsh, M.J. Kernan, and C. Kim. 2004. Two interdependent TRPV channel subunits, inactive and Nanchung, mediate hearing in Drosophila. *The Journal of neuroscience : the official journal of the Society for Neuroscience*. 24:9059-9066.

Gopfert, M.C., A.D. Humphris, J.T. Albert, D. Robert, and O. Hendrich. 2005. Power gain exhibited by motile mechanosensory neurons in Drosophila ears. *Proceedings of the National Academy of Sciences of the United States of America*. 102:325-330.

Han, Y.G., B.H. Kwok, and M.J. Kernan. 2003. Intraflagellar transport is required in Drosophila to differentiate sensory cilia but not sperm. *Current biology : CB*. 13:1679-1686.

He, M., R. Subramanian, F. Bangs, T. Omelchenko, K.F. Liem, Jr., T.M. Kapoor, and K.V. Anderson. 2014. The kinesin-4 protein Kif7 regulates mammalian Hedgehog signalling by organizing the cilium tip compartment. *Nature cell biology*. 16:663-672.

Ishikawa, H., T. Ide, T. Yagi, X. Jiang, M. Hirono, H. Sasaki, H. Yanagisawa, K.A. Wemmer, D.Y. Stainier, H. Qin, R. Kamiya, and W.F. Marshall. 2014.

TTC26/DYF13 is an intraflagellar transport protein required for transport of motility-related proteins into flagella. *eLife*. 3:e01566.

Jacobson, S.G., A.V. Cideciyan, I.V. Peshenko, A. Sumaroka, E.V. Olshevskaya, L. Cao, S.B. Schwartz, A.J. Roman, M.B. Olivares, S. Sadigh, K.W. Yau, E. Heon, E.M. Stone, and A.M. Dizhoor. 2013. Determining consequences of retinal membrane guanylyl cyclase (RetGC1) deficiency in human Leber congenital amaurosis en route to therapy: residual cone-photoreceptor vision correlates with biochemical properties of the mutants. *Human molecular genetics*. 22:168-183.

Jana, S.C., M. Girotra, and K. Ray. 2011. Heterotrimeric kinesin-II is necessary and sufficient to promote different stepwise assembly of morphologically distinct bipartite cilia in Drosophila antenna. *Molecular biology of the cell*. 22:769-781.

Jana, S.C., S. Mendonca, S. Werner, and M. Bettencourt-Dias. 2016. Methods to Study Centrosomes and Cilia in Drosophila. *Methods in molecular biology*. 1454:215-236.

Karpenahalli, M.R., A.N. Lupas, and J. Soding. 2007. TPRpred: a tool for prediction of TPR-, PPR- and SEL1-like repeats from protein sequences. *BMC bioinformatics*. 8:2.

Kolesova, H., M. Capek, B. Radochova, J. Janacek, and D. Sedmera. 2016. Comparison of different tissue clearing methods and 3D imaging techniques for visualization of GFP-expressing mouse embryos and embryonic hearts. *Histochemistry and cell biology*. 146:141-152.

Kuhn, M. 2016. Molecular Physiology of Membrane Guanylyl Cyclase Receptors. *Physiological reviews*. 96:751-804.

Lee, N., J. Park, Y.C. Bae, J.H. Lee, C.H. Kim, and S.J. Moon. 2018. Time-Lapse Live-Cell Imaging Reveals Dual Function of Oseg4, Drosophila WDR35, in Ciliary Protein Trafficking. *Molecules and cells*. 41:676-683.

Li, A., M. Saito, J.Z. Chuang, Y.Y. Tseng, C. Dedesma, K. Tomizawa, T. Kaitsuka, and C.H. Sung. 2011. Ciliary transition zone activation of phosphorylated Tctex-1 controls ciliary resorption, S-phase entry and fate of neural progenitors. *Nature cell biology*. 13:402-411.

Liu, X., K. Fujinami, K. Kuniyoshi, M. Kondo, S. Ueno, T. Hayashi, K. Mochizuki, S. Kameya, L. Yang, Y. Fujinami-Yokokawa, G. Arno, N. Pontikos, H. Sakuramoto, T. Kominami, H. Terasaki, S. Katagiri, K. Mizobuchi, N. Nakamura, K. Yoshitake, Y. Miyake, S. Li, T. Kurihara, K. Tsubota, T. Iwata, K. Tsunoda, and C. Japan Eye Genetics. 2020. Clinical and Genetic Characteristics of 15 Affected Patients From 12 Japanese Families with GUCY2D-Associated Retinal Disorder. *Translational vision science & technology*. 9:2.

Mangeol, P., B. Prevo, and E.J. Peterman. 2016. KymographClear and KymographDirect: two tools for the automated quantitative analysis of molecular and cellular dynamics using kymographs. *Molecular biology of the cell*. 27:1948-1957.

McGuire, S.E., P.T. Le, A.J. Osborn, K. Matsumoto, and R.L. Davis. 2003. Spatiotemporal rescue of memory dysfunction in Drosophila. *Science*. 302:1765-1768.

Morton, D.B. 2004. Atypical soluble guanylyl cyclases in Drosophila can function as molecular oxygen sensors. *The Journal of biological chemistry*. 279:50651-50653.

Mourao, A., S.T. Christensen, and E. Lorentzen. 2016. The intraflagellar transport machinery in ciliary signaling. *Current opinion in structural biology*. 41:98-108.

Nielsen, H. 2017. Predicting Secretory Proteins with SignalP. *Methods in molecular biology*. 1611:59-73.

Roca, A.I., A.E. Almada, and A.C. Abajian. 2008. ProfileGrids as a new visual representation of large multiple sequence alignments: a case study of the RecA protein family. *BMC bioinformatics*. 9:554.

Salehi Chaleshtori, A.R., M. Garshasbi, A. Salehi, and M. Noruzinia. 2020. The Identification and Stereochemistry Analysis of a Novel Mutation p.(D367Tfs*61) in the CYP1B1 Gene: A Case Report. *Journal of current ophthalmology*. 32:114-118.

Salvaterra, P.M., and T. Kitamoto. 2001. Drosophila cholinergic neurons and processes visualized with Gal4/UAS-GFP. *Brain research. Gene expression patterns*. 1:73-82.

Schindelin, J., I. Arganda-Carreras, E. Frise, V. Kaynig, M. Longair, T. Pietzsch, S. Preibisch, C. Rueden, S. Saalfeld, B. Schmid, J.Y. Tinevez, D.J. White, V. Hartenstein, K. Eliceiri, P. Tomancak, and A. Cardona. 2012. Fiji: an open-source platform for biological-image analysis. *Nature methods*. 9:676-682.

Tran, P.V., C.J. Haycraft, T.Y. Besschetnova, A. Turbe-Doan, R.W. Stottmann, B.J. Herron, A.L. Chesebro, H. Qiu, P.J. Scherz, J.V. Shah, B.K. Yoder, and D.R. Beier. 2008. THM1 negatively modulates mouse sonic hedgehog signal transduction and affects retrograde intraflagellar transport in cilia. *Nature genetics*. 40:403-410.

Tucker, C.L., V. Ramamurthy, A.L. Pina, M. Loyer, S. Dharmaraj, Y. Li, I.H. Maumenee, J.B. Hurley, and R.K. Koenekoop. 2004. Functional analyses of mutant recessive GUCY2D alleles identified in Leber congenital amaurosis patients: protein domain comparisons and dominant negative effects. *Molecular vision*. 10:297-303.

Wei, Q., Y. Zhang, Y. Li, Q. Zhang, K. Ling, and J. Hu. 2012. The BBSome controls IFT assembly and turnaround in cilia. *Nature cell biology*. 14:950-957.

Williams, C.L., J.C. McIntyre, S.R. Norris, P.M. Jenkins, L. Zhang, Q. Pei, K. Verhey, and J.R. Martens. 2014. Direct evidence for BBSome-associated intraflagellar transport reveals distinct properties of native mammalian cilia. *Nature communications*. 5:5813.

Wolff, A., B. de Nechaud, D. Chillet, H. Mazarguil, E. Desbruyeres, S. Audebert, B. Edde, F. Gros, and P. Denoulet. 1992. Distribution of glutamylated alpha and beta-tubulin in mouse tissues using a specific monoclonal antibody, GT335. *European journal of cell biology*. 59:425-432.

Yates, A., W. Akanni, M.R. Amode, D. Barrell, K. Billis, D. Carvalho-Silva, C. Cummins, P. Clapham, S. Fitzgerald, L. Gil, C.G. Giron, L. Gordon, T. Hourlier, S.E. Hunt, S.H. Janacek, N. Johnson, T. Juettemann, S. Keenan, I. Lavidas, F.J. Martin, T. Maurel, W. McLaren, D.N. Murphy, R. Nag, M. Nuhn, A. Parker, M. Patricio, M. Pignatelli, M. Rahtz, H.S. Riat, D. Sheppard, K. Taylor, A. Thormann, A. Vullo, S.P. Wilder, A. Zadissa, E. Birney, J. Harrow, M. Muffato, E. Perry, M. Ruffier, G. Spudich, S.J. Trevanion, F. Cunningham, B.L. Aken, D.R. Zerbino, and P. Flicek. 2016. Ensembl 2016. *Nucleic acids research*. 44:D710-716.

Ye, F., D.K. Breslow, E.F. Koslover, A.J. Spakowitz, W.J. Nelson, and M.V. Nachury. 2013. Single molecule imaging reveals a major role for diffusion in the exploration of ciliary space by signaling receptors. *eLife*. 2:e00654.

Ye, J., G. Coulouris, I. Zaretskaya, I. Cutcutache, S. Rozen, and T.L. Madden. 2012. Primer-BLAST: a tool to design target-specific primers for polymerase chain reaction. *BMC bioinformatics*. 13:134.

Zagel, P., and K.W. Koch. 2014. Dysfunction of outer segment guanylate cyclase caused by retinal disease related mutations. *Frontiers in molecular neuroscience*. 7:4.
